# Metabolomic markers of colorectal tumor with different clinicopathological features

**DOI:** 10.1101/2020.04.02.021899

**Authors:** Zhiping Long, Junde Zhou, Kun Xie, Zhen Wu, Huihui Yin, Volontovich Daria, Jingshen Tian, Nannan Zhang, Liangliang Li, Yashuang Zhao, Fan Wang, Maoqing Wang, Yunfu Cui

## Abstract

**Background:** Colorectal cancer (CRC) is the result of complex interactions between the tumor’s molecular profile and metabolites produced by its microenvironment. Despite recent studies identifying CRC molecular subtypes, a metabolic classification system is still lacking. We aimed to explore the distinct phenotypes and subtypes of CRC at the metabolic level.

**Methods:** We conducted an untargeted metabolomics analysis of 51 paired tumor tissues and adjacent mucosa using ultra-performance liquid chromatography/quadrupole time-of-flight mass spectrometry. Multivariate analysis including principal component analysis, orthogonal partial least squares discriminant analysis and heat maps, univariate analysis, and pathway analysis were used to identify potential metabolic phenotypes of CRC. Unsupervised consensus clustering were used to identify robust metabolic subtypes, and evaluated their clinical relevance.

**Results:** A total of 173 metabolites (including nucleotides, carbohydrates, free fatty acids, and choline) were identified between CRC tumor tissue and adjacent mucosa. We found that lipid metabolism was closely related to the occurrence and progression of CRC and CRC tissues could be divided into three subtypes, and statistically significant correlations between different subtypes and clinical prognosis were observed.

**Conclusions:** CRC tumor tissue exhibits distinct metabolic phenotypes. Metabolic differences between subtypes may provide a basis and direction for further clinical individualized treatment planning.

## INTRODUCTION

Colorectal cancer (CRC) is one of the leading causes of cancer-related death, both in China and worldwide. More than one million individuals develop CRC every year and most patients are diagnosed at advanced stages that correspond to poor prognosis (1). With the advances in the treatment of CRC over the past 20 years, median overall survival has been steadily increasing (2). Although the progress made thus far is encouraging, the existing treatment paradigm usually employs a ‘one-size-fits-all’ approach based on the histopathological diagnosis of CRC, which translates into demonstrable clinical benefit from any given chemotherapeutic regimen in only a small subset of treated patients (3).

It is now being increasingly realized that CRC is not a single disease entity, but a heterogeneous group of tumors, both at the intertumoral and intratumoral level (2). A major hallmark of CRC is its association with various types of etiological factors and its high heterogeneity in clinical presentation and underlying tumor biology (4). Consequently, most patients with CRC are refractory to treatment and have a dismal outcome. One of the essential requirements to improve their outcome is to provide biomarkers that are capable of accurately defining homogenous molecular subtypes; each displays unique tumor biology linked to potentially druggable driver genes to implement rational treatment choices (5).

Nowadays, tumor genomic profiling is routinely used to classify tumor types, identify driver or germline mutations, perform prognostic assessments, and make therapeutic decisions (6, 7). However, the notable heterogeneity of genomes in cancer tissues makes it difficult to determine the underlying causes or ascertain the optimal treatment. Furthermore, the elevated number of mutations and multiple combinations of tumor suppressors and oncogenes make individualized tumor classification or customized therapy almost impossible (8). Metabolomics is a rapidly growing field of study that endeavors to measure the complete set of metabolites (generally considered to be the intermediates and products of cellular metabolism less than 1 kDa in size) within a biological sample (that is, the metabolome) to achieve a global view of the state of the system (9). In general, multiple biochemical pathways are affected, owing to the fact that as cancer progresses, multiple defects in biochemical pathways arise as cancer subverts normal metabolism in an effort to survive (10). Furthermore, the metabolic requirements of cancer cells are different from those of most normal differentiated cells, exhibiting different metabolic phenotypes (11). Using metabolomics to identify the specific metabolic subtype of a particular tumor would enable better customization or informed adjustment of cancer therapies (12).

To present, metabolomics-based CRC phenotypic research and molecular typing have been rarely described, and little is known about how changes in metabolite levels relate to the characteristics of tumor tissue. In this study, we described a metabolomics analysis of CRC tissue samples from a group of CRC patients with different clinicopathological features. We aimed to analyze the differential metabolism of tumor tissues with different clinicopathological features, and to explore molecular typing methods for CRC based on metabolomics markers.

## METHODS

### Study design and subject recruitment

We designed a self-control study to detect the differential metabolites between tumor tissue and adjacent non-malignant mucosa tissue. Fifty-one pairs of tissue were obtained from surgical resection of CRC patients.

All patients were diagnosed and recruited at the Third Affiliated Hospital of Harbin Medical University. Any patients with neuroendocrine carcinoma, malignant melanoma, non-Hodgkin’s lymphoma, gastrointestinal stromal tumors, and Lynch syndrome CRC were excluded. Only newly diagnosed histopathologically confirmed cases were retained. Tissue sampling included the deepest infiltration of the tumor and the adjacent non-malignant mucosa tissues. All tissues were immediately soaked in formaldehyde solution until use.

All procedures performed in studies involving human participants were in accordance with the ethical standards of the Human Research and Ethics Committee of Harbin Medical University and with the 1964 Helsinki declaration and its later amendments or comparable ethical standards. Informed consent was obtained from all individual participants included in the study.

### Metabolic profiling

A detailed description of the experimental protocol of metabolic profiling analysis by UPLC/Q-TOF-MS/MS and the data processing, multivariate and univariate analysis of metabolites, as well as identification of differential metabolites, are provided in the Supplementary Materials.

### Pathway analysis

Using an accurate m/z search under 50 ppm, metabolites from positive and negative ionization were matched in Mummichog software, which included metabolites from KEGG and other databases. Mummichog software (version 1.0.9) was used to further test pathway enrichment patterns using permutations, and to compute the probability for each pathway (13).

### Metabolic clustering

Consensus clustering (cCluster; hierarchical clustering; Pearson distance; complete linkage; 1000 resampling iteration) and unsupervised hierarchical clustering were performed to define subtypes of CRC tumor tissue samples (14, 15). Heatmaps were generated using the Complex Heatmap package in R to determine the relationship among samples or cCluster-defined subgroups (16).

### Clinical relevance analysis of metabolic subtypes

We assessed whether the metabolic subtypes had significant associations with overall survival. The R packages “survival” and “survminer” were used to perform the overall survival analysis and to produce Kaplan-Meier survival plots. A log-rank test was used to assess the significance (*P*<0.05). We further assessed whether the metabolic subtypes remained significantly associated with overall survival after adjusting for age, sex, TNM staging, postoperative chemotherapy, and immunotherapy as covariates in the Cox model.

## RESULTS

### Metabolic profiling of 51 pairs of tumor tissue and adjacent mucosa tissue

To identify the differential metabolites of CRC, the metabolomes of tumor tissues were compared with that of matched adjacent mucosa. Supplementary Table 1 shows the demographic characteristics and clinicopathological features of 51 CRC patients. Mass spectrometry detected 4,526 and 4,765 variables in negative electrospray ionization (ESI−) and positive electrospray ionization (ESI+), respectively. The statistical variation was visualized using OPLS-DA (Figure 1). OPLS-DA revealed the metabolites that significantly contributed to this variation and their relationship with one state versus another. A total of 373 metabolites (296 higher and 77 lower) were identified with the criteria of Variable important for the projection (VIP) score greater than 1.5 and *P* values of less than 0.05 in the false detection rate (FDR)-corrected Mann-Whitney U tests, which displayed differential abundance between tumor and adjacent mucosa samples (Supplementary Figure 1). The Human Metabolome Database (http://www.hmdb.ca/) mass search feature was used as to aid metabolite identification. A total of 173 metabolites were identified as shown in Supplementary Table 2. Interestingly, nucleotides, carbohydrates, free fatty acids, and choline were overrepresented and highly abundant in tumors, such as D-ribulose 5-phosphate, D-glucose, xylulose 5-phosphate, 3’-AMP, hypoxanthine, palmitoleic acid, and cytidine monophosphate (Supplementary Table 2).

**Figure 1.**
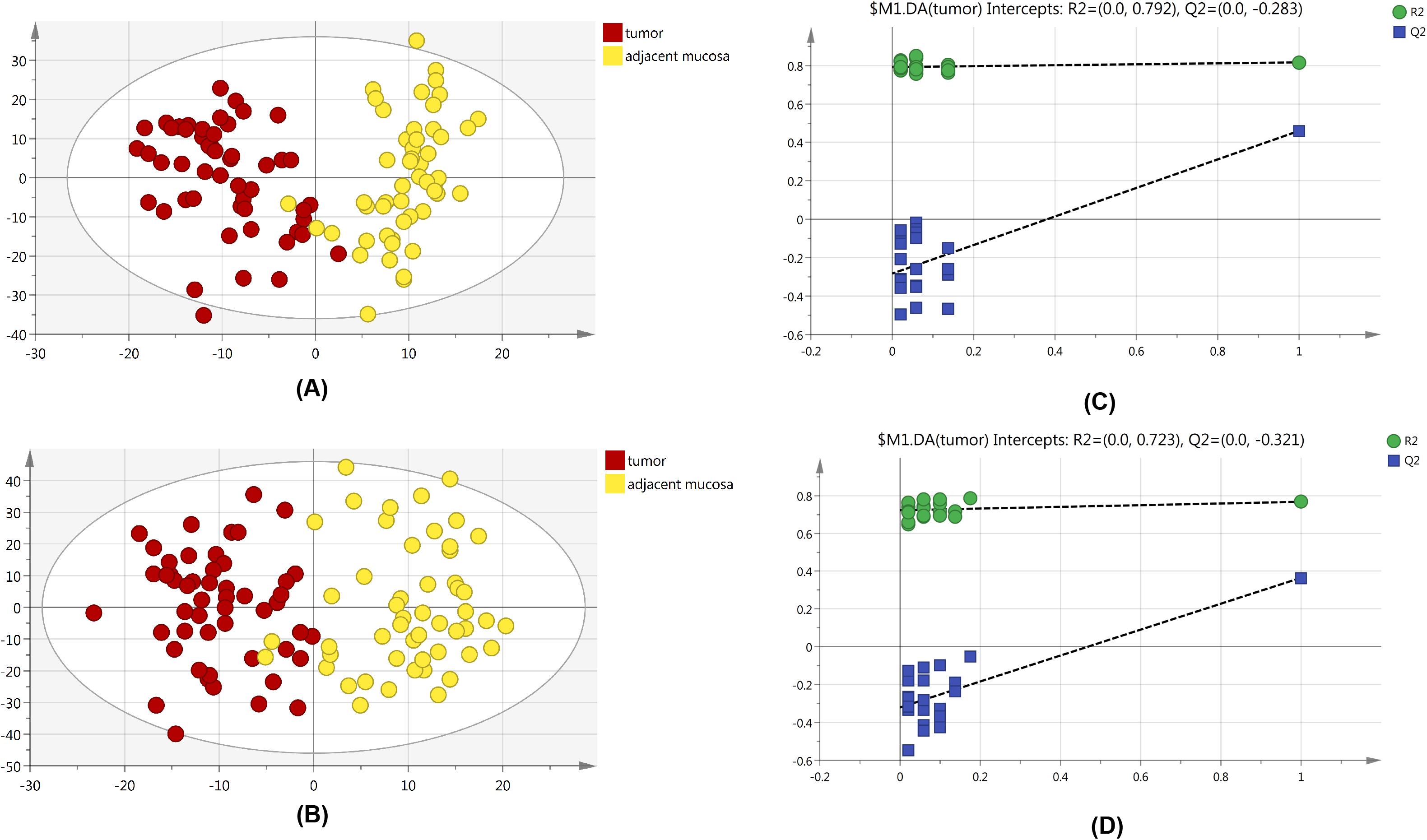
OPLS-DA score plots of tissue from colorectal cancer patients and permutation test results of OPLS-DA model. (A) OPLS-DA score plot in ESI^−^ model, R^2^X=0.218, R^2^Y=0.817, Q^2^=0.459. (B) OPLS-DA score plot in ESI^+^ model, R^2^X=0.246, R^2^Y=0.768, Q^2^=0.364. The R^2^Y value represents the goodness of fit of the model. The Q^2^ value represents the predictability of the model. (C) Permutation test result of the OPLS-DA model in ESI^−^ model.; (D) Permutation test result of the OPLS-DA model in ESI^+^ model.

### Metabolic landscape of CRC tumors

Pathway analysis was performed to systematically investigate the metabolic alterations associated with CRC pathogenesis. Mummichog software, a pathway tool designed for untargeted metabolomics data [13], was used to evaluate the significant metabolic pathways utilizing metabolites that were present at differential abundance between CRC tissues and adjacent mucosa. The results of Mummichog analysis are shown in Supplementary Table 3; interestingly, among the 34 metabolic pathways, most were involved in lipid metabolism (n=7) and glycan biosynthesis and metabolism (n=8). Other metabolic pathways included glycolysis/gluconeogenesis, pentose phosphate pathway, and tryptophan metabolism.

### Metabolic changes upon CRC cancer progression

Difference stage-distributed CRC samples allowed us to investigate the association between metabolic shifts and CRC progression. As shown in Figure 2, using American Joint Committee on Cancer (AJCC) clinical staging in both the positive and negative mode, each stage segregated well in OPLS-DA. Furthermore, as shown in the heatmaps in Figure 4, it was possible to identify metabolic features that distinguished the various stages of CRC patients. There were 94 metabolites exhibiting statistically significant differential abundance between early- (I, II) and late-stage (III, IV) tumors (VIP >1.5 and *P*<0.01), and a total of 48 metabolites were identified (Supplementary Table 3). Especially, most lipid metabolites showed an increase in late-stage tumors, while dipeptides also showed a decrease in late-stage tumors (Supplementary Figure 2). The results of pathway analysis by Mummichog software indicated that significant features are enriched for pathways involved in lipid metabolism (Supplementary Table 4).

**Figure 2.**
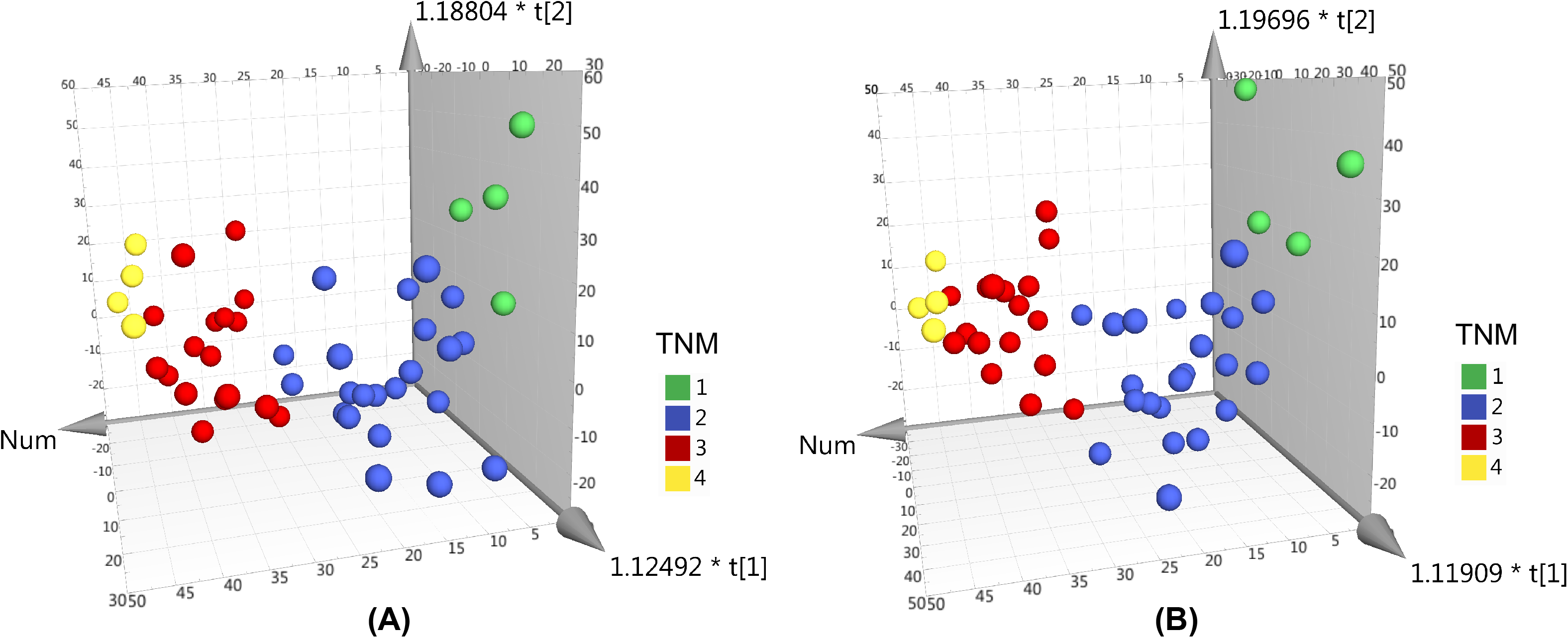
OPLS-DA 3D scores plots of tissue from various stages CRC patients. (A) OPLS-DA score plot in ESI^−^ model, R^2^X=0.199, R^2^Y=0.34, Q^2^=0.042. (B) OPLS-DA score plot in ESI^+^ model, R^2^X=0.2, R^2^Y=0.335, Q^2^=0.0599.

### Metabolic alterations of CRC pathologic characteristics

We also sought to determine whether we could identify the differences in metabolic features among various histopathological classifications of CRC. Figure 3 shows the separation of adenocarcinoma and non-adenocarcinoma CRCs using OPLS-DA. Forty-three metabolites exhibited statistically significant differential abundance between adenocarcinoma and non-adenocarcinoma tumors (VIP >1.5 and *P*<0.01). Furthermore, a total of 26 metabolites were identified (Supplementary Table 5) and almost all these 26 metabolites were lipids. Mummichog indicated that pathways involved in lipid metabolism were also significantly enriched (Supplementary Table 5).

**Figure 3.**
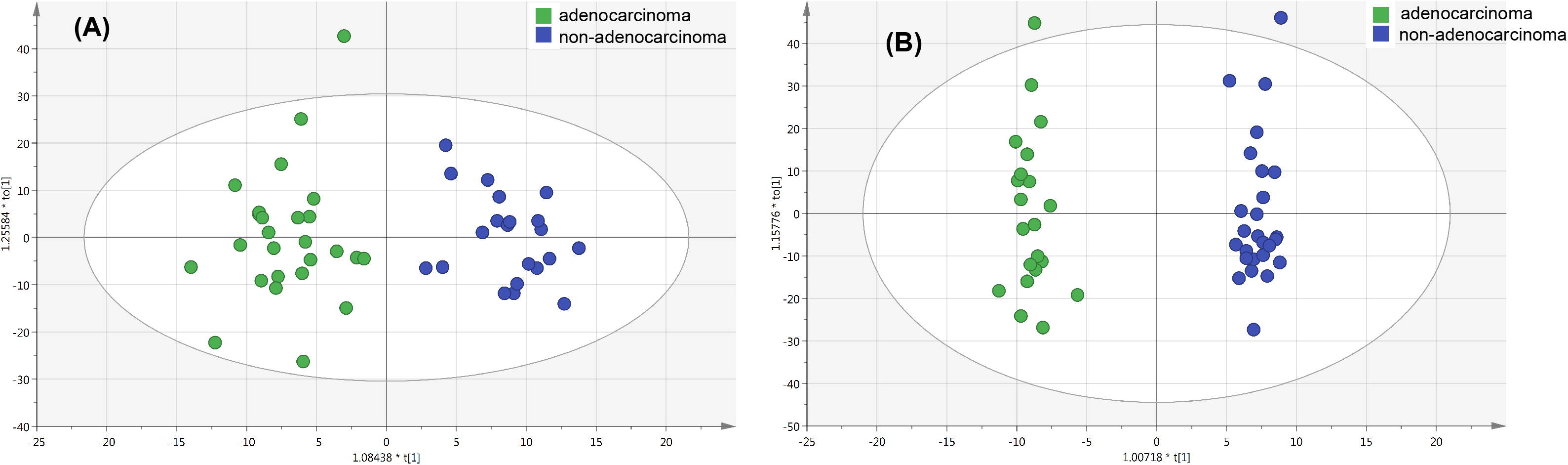
OPLS-DA score plots of tissue from various histopathologic classification CRC patients. (A) OPLS-DA score plot in ESI^−^ model, R^2^X=0.088, R^2^Y=0.876, Q^2^=0.128. (B) OPLS-DA score plot in ESI^+^ model, R^2^X=0.229, R^2^Y=0.984, Q^2^=−0.026.

### Unsupervised clustering reveals three metabolic clusters (mClusters) with prognostic value

The results of cCluster showed that CRC tumor samples can be partitioned into clusters with distinct metabolic phenotypes using the differential metabolites among tumor and adjacent mucosa samples. cCluster revealed three major subtypes of CRC according to consensus distributions and the corresponding consensus matrices (Supplementary Figure 3). The CRC subtypes defined by cCluster can be visualized through unsupervised hierarchical clustering (Supplementary Figure 3). This analysis revealed unique subtypes of CRC cases with distinct metabolic patterns that were independent of known clinicopathological features (Figure 4). For each metabolite cluster (mCluster), the clinical stages at presentation are summarized in Supplementary Figure 4. mCluster 1 had the highest percentage (66.7%) of early-stage (I & II) tumors and was characterized by the low abundance of carbohydrates, nucleotide metabolites, dipeptides, and lipids; mCluster 2 had the highest percentage (51.9%) of late-stage (III & IV) tumors and displayed medium levels of all metabolites; mCluster 3, characterized by the highest abundance of carbohydrates, nucleotide metabolites, dipeptides, and lipids, accounted for 62.5% early-stage tumors (Figure 4 and Supplementary Figure 4). Additionally, we further determined the correlations of mClusters with patients’ overall survival, since survival represents a critical clinical index of tumor aggressiveness. As shown in Figure 5, the result did not reach statistical significance, likely due to the relatively small number of events during follow-up (log-rank *P*=0.099). However, after mCluster 1 and 3 were combined, cases defined as mCluster 2 showed poor survival (log-rank *P*=0.032). We obtained similar results using Cox regression models adjusting by age, sex, TNM staging, and postoperative chemotherapy and immunotherapy (*P*=0.027) (Supplementary Figure 5).

**Figure 4.**
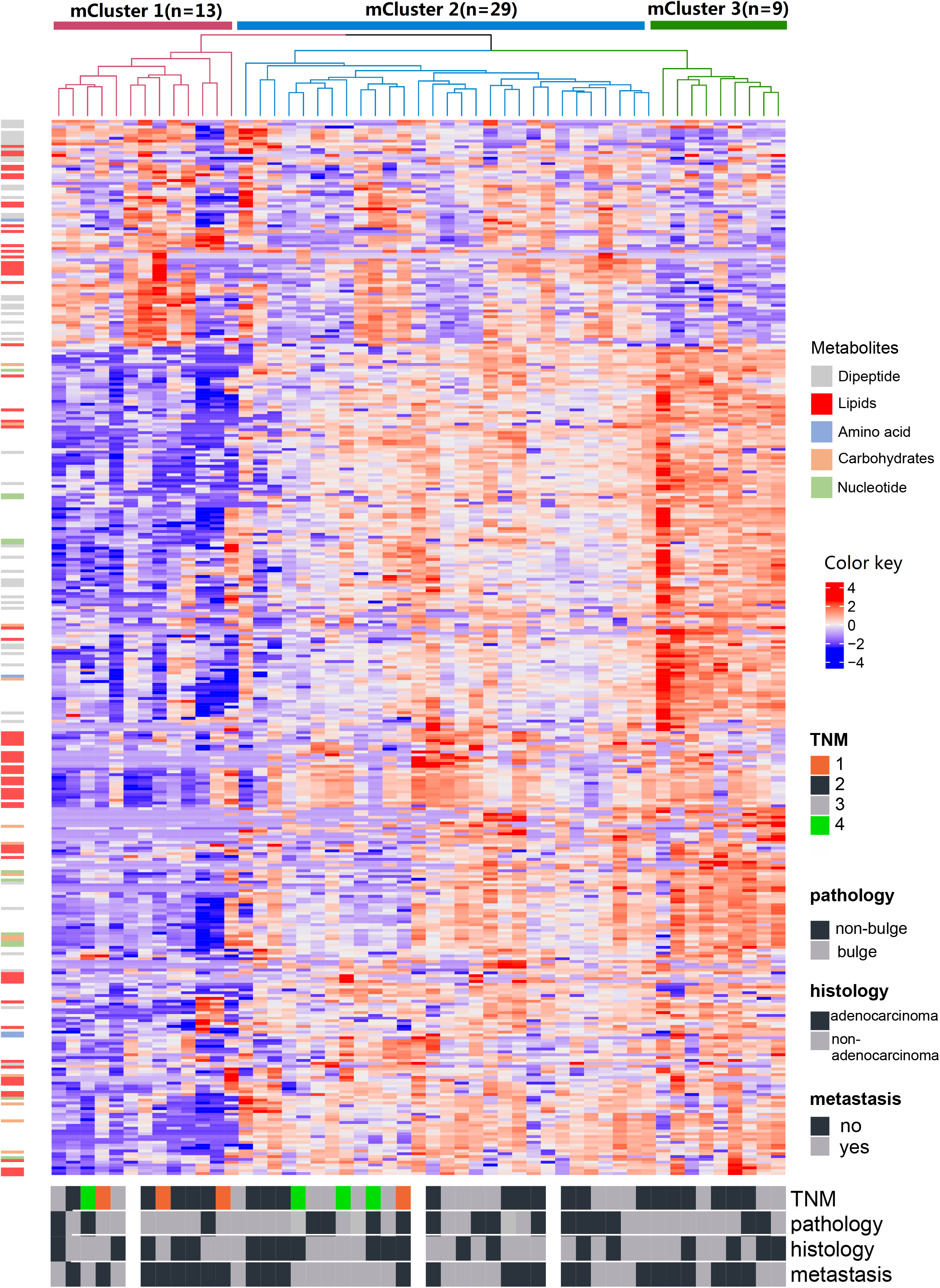
Identification of CRC metabolite-based tumor subtypes. A heatmap of CRC subtypes is shown based on consensus clustering. The x-axis represents CRC subtype consensus clusters. CRC samples are represented in columns, grouped by the dendrogram into three main clusters, and metabolites (n=373) are represented in rows. Clinical data of the samples are included below the heatmap.

**Figure 5.**
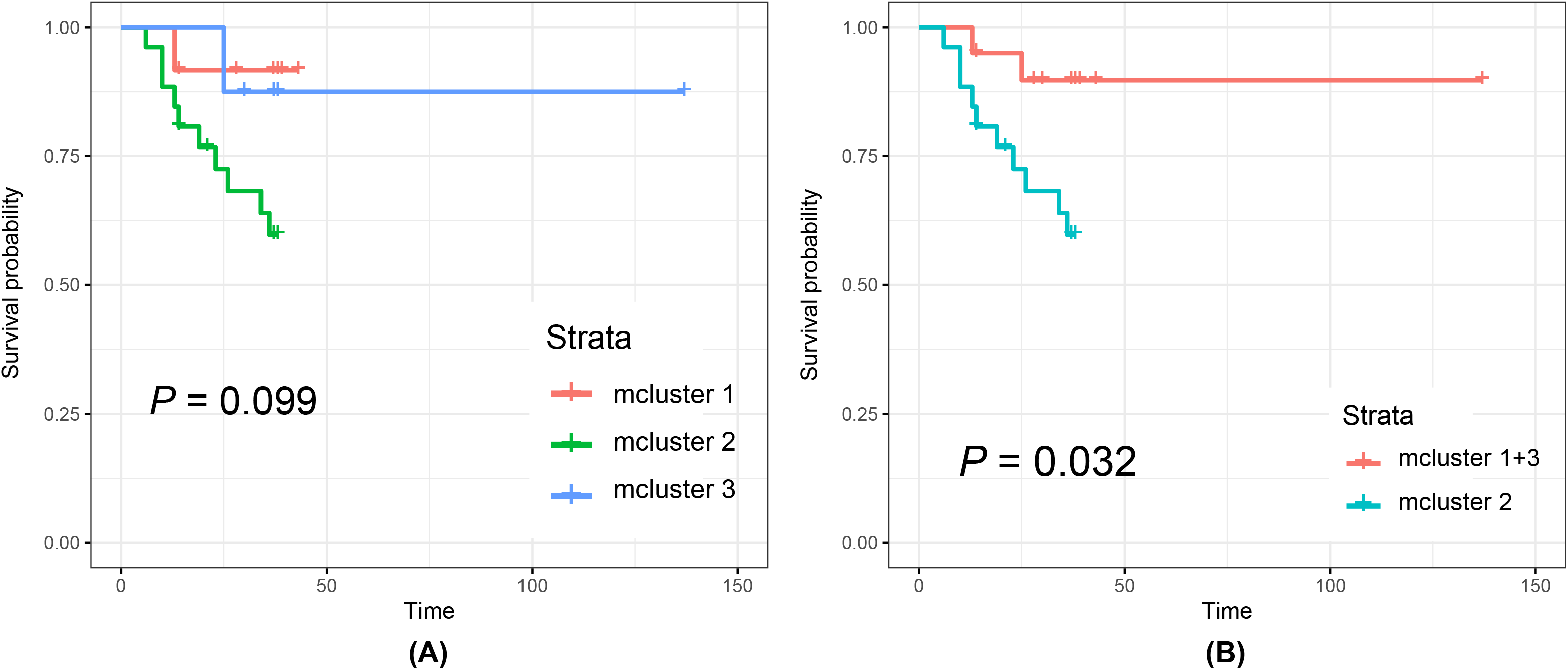
Kaplan-Meier curves of cancer-specific survival of individual mclusters.

## DISCUSSION

Metabolomics analysis of CRC can not only distinguish tumor tissue from adjacent mucosa, but can also discriminate CRC patients with different clinicopathological features. Metabolic profiling allows a more comprehensive understanding of CRC phenotyping, and we can define molecular subtypes of CRC based on metabolomics. The results of our study have shown that through the high-throughput metabolomics analysis using UPLC/Q-TOF MS mass spectrometry platform, many significant differential metabolites can be found between CRC tumor tissues and adjacent mucosa. Based on unsupervised and supervised results, there were differential metabolism patterns between early- and late-stage CRCs, and CRC tumor tissues could be divided into three metabolic subtypes with differential prognosis.

Tian et al. analyzed the metabonomic signatures of 50 human CRC tissues and their adjacent non-involved tissues (ANIT) using high-resolution magic-angle spinning (HRMAS) 1H NMR spectroscopy together with the fatty acid compositions of these tissues using GC-FID/MS(17). In this study, metabonomic phenotypes of CRC tissues differed significantly from that of ANIT in energy metabolism, membrane biosynthesis and degradation, and osmotic regulation together with the metabolism of proteins and nucleotides. Diverse metabolic pathways including N-glycan biosynthesis and degradation, linoleate metabolism, leukotriene metabolism, butanoate metabolism, glycosphingolipid biosynthesis, drug metabolism-cytochrome P450 and vitamin B5-CoA biosynthesis from pantothenate significantly differed between tumor and normal tissues. The UPLC/Q-TOF MS-based metabolomics approach of this study provided additional information that complements our current understanding of the metabolomic characteristics between CRC tumor tissues and adjacent mucosa.

The Warburg effect is a known feature of cancer metabolism that describes maintenance of a high aerobic glycolysis rate and high levels of glucose uptake and lactate production during tumor growth (18, 19). Our findings are consistent with the Warburg effect. The difference in energy metabolism can be clearly observed between CRC tumor tissues and adjacent mucosa. Compared with adjacent mucosa, carbohydrates in colorectal cancer tissues were significantly increased and the pentose phosphate pathway and glycolysis/gluconeogenesis pathways were identified. In cancer metabolism, glycolysis is the preferred pathway to produce metabolic intermediates used to support cell proliferation during de novo biosynthesis (20), which can lead to higher levels of free fatty acids (FFA) and nucleic acid-related metabolites. In our current study, higher levels of nucleotides, palmitoleic acid, and hypoxanthine were observed in tumor tissues. Nucleotides are critical components of DNA and RNA structures, and disorders in their biosynthesis have profound effects on cell physiology, which may lead to tumor transformation in cells (21). CRC tumor tissues showed higher levels of choline metabolites such as choline, PC, and PE than adjacent mucosa, which have also been reported in other malignancies (22–24).

Glycosylation changes are some of the most common post-translational modifications of proteins and are considered markers of cancer. N-glycans can regulate cell migration, cell adhesion, cell signaling, proliferation, and metastasis. Many carbohydrate-mediated cellular mechanisms, including those important for tumor progression, are regulated by N-glycans (25). Stephanie et al. compared the glycosylation profiles of tumor tissues and corresponding control tissues in 13 colorectal cancer patients (26). Multivariate data analysis showed significant differences in glycosphingolipids between tumors and corresponding adjacent tissues using MALDI-TOF(/TOF)-MS and 2-dimensional LC-MS/MS; the main changes included elevated fucosylation, reduced acetylation and sulfation, and reduced expression of globular glycans, as well as disialyl gangliosides. In our study, seven metabolic pathways were identified as being involved in the biosynthesis and metabolism of glycans, including biosynthesis of N-glycans, degradation of N-glycans, and metabolism and biosynthesis of glycosphingolipids, confirming the changes in characteristic tumor-associated glycosylation.

To date, there have been few studies analyzing the differences in the metabolism of CRC with different clinicopathological features. In this study, it was reported for the first time that the early tumors of CRC have higher abundance of dipeptide characteristics. A large increase in dipeptides may be produced through protein degradation/reutilization processes, such as lysosomal degradation, phagocytosis, endocytosis, pinocytosis, and autophagy (27–30). Brauns et al. have shown that cyclic dipeptides, especially those containing proline, have important biological activities. Their results indicated that phenylalanine–proline inhibits the proliferation of HT-29, MCF-7, and HeLa cells, as well as inducing apoptosis in HT-29 colon cancer cells, which has potential anti-tumor activity (31).

Higher levels of lipid metabolites observed in the current study in advanced CRC tissues have been reported in other studies (17, 32). Results of the Mummichog software pathway analysis showed that most pathways are lipid metabolism-related, consistent with previous studies by Zhang et al. and Tian et al. (33, 34). Abnormal lipid metabolism is a metabolic marker of cancer cells (35, 36), and many studies have reported that by activating the exogenous (or dietary) lipid and lipoprotein uptake or by enhancing the reticular fat from the cytosol acetyl-CoA Biosynthesis of cholesterol and cholesterol, highly proliferative cancer cells have strong lipid and cholesterol affinities (36). Changes in lipid metabolism in CRC tumor tissues suggest enhanced lipogenesis is one of the most important features in CRC tumor tissues (37). Recent studies have also found that tumor tissue can use fatty acids and lipolytic pathways to obtain fatty acids to promote tumor cell proliferation (38).

We further observed that the metabolic differences between adenocarcinoma and non-adenocarcinoma CRCs were mainly related to lipid metabolism. Lipid metabolism is regulated by complex signaling networks in CRC tumor cells, which are closely related to cell growth, proliferation, differentiation, survival, and apoptosis (39). Several studies have indicated that some fatty acid metabolism pathways are associated with the development and progression of colorectal adenocarcinoma (40, 41). Beatriz et al. also showed that changes in fatty acid metabolism are a crucial factor in the progression from colorectal adenoma to adenocarcinoma (42). Although our results are consistent with previous studies, there have been no studies on the metabolic differences of adenocarcinoma and non-adenocarcinoma thus far.

Our results, for the first time, showed that CRC could be divided into three subtypes at the metabolomics level, and that the subtypes were significantly associated with prognosis. Lipids, nucleotides, and carbohydrates have important roles in the biology of a subset of tumors. The differences in these metabolite levels between subtypes may point to different pathophysiological mechanisms for the development and progression of CRC. Understanding the pathogenesis of CRC is critical to developing personalized treatment strategies. CRC is currently still classified according to the TNM or TNM clinical staging system, an approach that is primarily focused on the surgical treatment of CRC but not personalized systemic treatments. As every CRC covers a specific, heterogeneous metabolic profile, the question rises if metabolomics (and other ‘omics’) approaches could become the new standard in adequately categorizing CRC on a molecular basis. This molecular classification could offer patients a personalized therapy schedule, depending on the type of molecular defects that their colorectal tumor has acquired.

For example, many anticancer drugs are based on lipid metabolism, such as irinotecan, which can affect the accumulation of ceramide by inducing ceramide synthase to catalyze ceramide synthesis or by activating sphingomyelinase to catalyze the degradation of sphingomyelin (43, 44). At the same time, the use of drugs is also dependent on the sensitivity and intrinsic drug resistance of cancer cells. Studies have shown that omega-3 polyunsaturated fatty acids can improve the efficacy of chemotherapy and radiotherapy. Omega-3 fatty acids also reduce CD133+ colon cancer stem cell-like cells markers and increase sensitivity to chemotherapy (45). A eicosapentaenoic acid-free fatty acid(EPA-FFA) phase II double-blind, placebo-controlled trial of patients undergoing liver resection for CRC liver metastases showed that EPA-FFA treatment is anti-angiogenic, safe, and well tolerated (46). Backshall et al. evaluate the effect of pretreatment serum metabolic profiles generated by 1H NMR spectroscopy on toxicity in patients with inoperable colorectal cancer receiving single agent capecitabine (47). Their study suggests that metabolic profiles can delineate subpopulations susceptible to adverse events and have a potential role in the assessment of treatment viability for cancer patients prior to commencing chemotherapy.

This study still has some limitations. Our study is based on a relatively small sample of colorectal cancer patients in northeastern China. Moreover, the UPLC/Q-TOF MS metabolomics platform used in the study was used in isolation and some metabolites may not have been detected. Therefore, confirmation is necessary based on multiple populations and platforms.

In summary, our metabolomic study indicates that CRC tumor tissue exhibits distinct metabolic phenotypes. Metabolomics provides a new window into the study of CRC phenotypes and molecular typing as CRC can be divided into three subtypes at the metabolic level. When integrated with other platforms, we can provide a more comprehensive explanation of the complex biology associated with CRC and malignant transformation. A deeper understanding of abnormal metabolism will provide a framework for the design and implementation of personalized approaches to CRC treatment through metabolic regulation.

## Abbreviations

CRC: colorectal cancer
ESI−: negative electrospray ionization
ESI+: positive electrospray ionization
VIP: variable important for the projection
mClusters: metabolic clusters
FFA: free fatty acids
ANIT: adjacent non-involved tissues

## Acknowledgments

This work was supported by grants from National Natural Science Foundation of China (81773503, 81573147), Scientific Research Foundation for the Returned Overseas Scholars of Heilongjiang Province (LC2018033), and Dr. Wu Lien-teh Science Foundation of Harbin Medical University (WLD-QN1106).

