## Supplementary Material for "Metabolomic markers of colorectal tumor with different clinicopathological features"

### I. Supplementary Methods

#### 1.1 Experimental protocol of metabolic profiling analysis for tissue by UPLC/Q-TOF-MS/MS

*Chemicals and reagents* Acetonitrile and methanol (HPLC grade) were purchased from Honeywell Burdick & Jackson (Muskegon, MI). Formic acid was purchased from Beijing Reagent Company (Beijing, China). All chemicals and regents were of HPLC grade available from commercial sources. Ultrapure water was prepared by an ultra-clear system (PURELAB Ultra; Veolia Water Solutions & Technologies, France).

*Tissue sample preparation* About 100 mg of tissue soaked in formaldehyde solution was placed in a mortar containing Liquid nitrogen and fully grounded. After the well-ground tissue was completely dissolved with 2 ml of methanol, all metabolites in the tissue were extracted and all proteins were precipitated. The solution obtained from step 1 was transferred to a 2 ml centrifuge tube, then was vortexed for 1 min; and centrifuged at 12,000 rpm for 10 min at 4°C. The supernatant was put in another 2 ml centrifuge tube and was dried with nitrogen. The substance dried by nitrogen in step 3 was dissolved with methanol (1:1) and the tissue soaking solution (300-400ul). Vortexed for 1 min and standing for 5 min, then centrifuged at 12,000 rpm for 10 min at 4°C. The supernatant was finally removed and transferred to auto sampler vial for metabolomics analysis by ULPLC/Q-TOF MSMS. A quality control sample (QC) was prepared by mixing aliquots from all supernatant samples (10 μL from each sample).

*UPLC/Q-TOF-MS/MS analysis* Chromatographic separation was performed on a 1.7-μm BEH C18 column [ACQUITY (HSS); Waters Corp., Milford, MA, USA; 2.1

mm × 100 mm] equipped with a UPLC system (ACQUITY UPLC; Waters Corp., USA). The temperatures of the column and autosampler were maintained at 35°C and 4°C, respectively. A sample (2 μl) of the preprocessed plasma was injected onto the column at a flow rate of 0.35 ml/min. The mobile phase consisted of water containing 0.1% formic acid waters (solution A) and acetonitrile (solution B). The elution gradient was as follows: 2% B for 0.5 min; 2% to 20% B over 0.5 to 6.0 min; 20% to 35% B over 6.0 to 7.0 min; 35% to 70% B over 7.0 to 9.0 min; 70% to 98% B over 9.0 to 10.5 min; 98% B for 2.0 min; and then a return to 2% B for 6.0 min. Once the initial settings had been established, the column was equilibrated for 2.0 min. Acetonitrile was run every fifth sample as a blank solution and the plasma samples in the two analysis batches were injected alternately as five cases and five control samples.

Q-TOF MS/MS was performed with a mass spectrometer (Micromass Q-TOF mass spectrometer; Waters Corp., Manchester, UK) using an electrospray ionization (ESI) interface operated in both ion modes ($ESI^-$ and $ESI^+$). The analytical parameters were as follows: capillary voltage, 2800 V in $ESI^-$ or 3000 in $ESI^+$; sample cone voltage, 35 V; collision energy, 6 eV; source temperature, 110°C; desolvation gas (nitrogen) flow, 650 L/h; desolvation temperature, 320°C; cone gas (nitrogen) flow, 50 L/h; collision gas, argon; and MCP detector voltage, 2400 V. The Q-TOF mass acquisition rate was set at 0.4 s, with an interscan delay of 0.1 s. The scan mass range was from 50 to 1000 m/z. The data were collected in centroid mode, using the lock spray to ensure accuracy and reproducibility. A concentration of 200 pg/ml leucine–enkephalin was used as lock mass (m/z 554.2615) in $ESI^-$ and (m/z 556.2771) in $ESI^+$. The lock spray frequency was set at 10 s, and the lock mass data were averaged over 10 scans for correction. The MS/MS spectra of metabolites were obtained by UPLC-MS/MS.

#### 1.2 Data processing, multivariate and univariate analysis of metabolites, and identification of differential metabolites

*Data processing* The raw data were imported into MarkerLynx software incorporated in Masslynx software (version 4.1 SCN714). MarkerLynx ApexTrack peak integration was used for peak detection and alignment. The ApexTrack peak parameters were set as follows: peak width at 5% height, 1 s, and peak-to-peak baseline noise (calculated automatically). Collection parameters were set as follows: retention time (RT) range 0.5–10.5 min, mass range 50–1000 Da, mass tolerance, 0.05 Da; RT tolerance, 0.1 min; minimum intensity, 80; noise elimination level, 6.0; and deisotope data, yes. After being recognized and aligned, the intensity of each ion was normalized to the summed total ion intensity of each chromatogram. The data-reduction process was handled in accordance with the “80% rule.”

*Multivariate analysis of metabolites* A matrix of samples against variables was generated and transferred to SIMCA-P version 13.0 software (Umetrics, Umeå, Sweden) for multivariate analysis. Principal component analysis (PCA) was first performed to check the outliers and the separation tendency. Orthogonal projections to latent structures discriminant analysis (OPLS-DA) was further applied to visualize the maximal difference between cases and controls. A default sevenfold (leave one-seventh of samples out) cross-validation procedure was used to assess the robustness of the models. Furthermore, permutation tests calculated by 100 randomizations were performed to avoid the overfitting of supervised OPLS-DA models. The variable importance in projection (VIP) values of all peaks from the OPLS-DA model was taken as a coefficient for peak selection.

*Univariate analysis of metabolites* The false detection rate (FDR)-corrected Mann-Whitney U tests was applied for the selection and validation of differential variables. The metabolites with statistical significance in both multivariate and univariate analyses (VIP >1.5 and $P$ <0.05) were identified.

*Differential metabolite identification* Metabolite annotation was performed by comparing the exact m/z values and MS/MS spectra with those in free online databases Human Metabolome Database (HMDB, http://www.hmdb.ca/). If the potential MS/MS spectra were not available in online databases, the MassFragment application manager (MassLynx version 4.1, Waters) was applied to facilitate the MS/MS fragment ion analysis process via chemically intelligent peak-matching algorithms. Differential metabolites were finally confirmed using standard compounds based on both retention time and MS/MS spectra.

### II. Supplementary Tables

**Supplementary Table 1. Demographic and pathological characteristics of fifty-one colorectal cancer patients.**

**Supplementary Table 2. Colorectal cancer metabolites identified from the paired-tissue metabolomics study.**

**Supplementary Table 3. Pathway analysis of 373 positive and negative ion by Mummichog.**

**Supplementary Table 4. Fifty identifiable metabolites differential abundance between early (I, II) and late-stage (III, IV) tumors.**

**Supplementary Table 5. Pathway analysis of 94 positive and negative ion by Mummichog**

**Supplementary Table 6. Twenty-six identifiable metabolites differential abundance between adenocarcinoma and non-adenocarcinoma tumors.**

**Supplementary Table 7. Pathway analysis of 43 positive and negative ion by Mummichog.**

### III. Supplementary Figures

**Supplementary Figure 1. Heatmap visualization constructed based on 373 differential variables.** Note: Rows, samples; columns, differential variables. Color key indicates metabolite expression value: dark blue, lowest; dark red, highest.

**Supplementary Figure 2. Heatmap visualization constructed based on 94 differential variables.** Note: Rows, samples; columns, differential variables. Color key indicates metabolite expression value: dark blue, lowest; dark red, highest.

**Supplementary Figure 3. Results of consensus clustering of 373 differential variables: Identification of four metabolomic subtypes.** (A) Consensus clustering matrix of 51 CRC samples for k=2 to k=6. (B) Consensus clustering CDF for k=2 to k=6. (C) The corresponding relative change in area under the cumulative distribution function (CDF) curves when cluster number changed from k to k+1. The range of k changed from 2 to 6 and the optimal k=3.

**Supplementary Figure 4. Clinical stages at sample collection and the eventual metastasis of each individual metabolic cluster.**

**Supplementary Figure 5. Cox analysis for the survival of patients with different mclusters**

**Supplementary Table-1. Demographic and pathological characteristics of fifty-one colorectal cancer patients.**

| No. | Age | Gender | Location | Lymphatic metastasis | Distal metastasis | TNM staging | Histological type |
|---|---|---|---|---|---|---|---|
| 1 | 65 | Male | Spleen area | 1 | 1 | 2 | Non-adenocarcinoma |
| 2 | 45 | Male | Rectum | / | / | / | / |
| 3 | 79 | Male | Rectum | 1 | 1 | 2 | Adenocarcinoma |
| 4 | 58 | Female | Rectum | 1 | 1 | 1 | Adenocarcinoma |
| 5 | 54 | Male | Sigmoid colon | 1 | 1 | 2 | Non-adenocarcinoma |
| 6 | 59 | Male | Liver area | 1 | 1 | 2 | Non-adenocarcinoma |
| 7 | 59 | Female | Rectum | 2 | 1 | 3 | Adenocarcinoma |
| 8 | 45 | Male | Ascending colon | 2 | 1 | 3 | Adenocarcinoma |
| 9 | 62 | Female | Rectum | 2 | 1 | 3 | Non-adenocarcinoma |
| 10 | 71 | Female | Cecum | 1 | 1 | 2 | Non-adenocarcinoma |
| 11 | 37 | Male | Sigmoid colon | 1 | 2 | 4 | Adenocarcinoma |
| 12 | 22 | Female | Transverse colon | 1 | 2 | 4 | Non-adenocarcinoma |
| 13 | 56 | Male | Descending colon | 1 | 1 | 2 | Non-adenocarcinoma |
| 14 | 43 | Female | Rectum | 2 | 1 | 3 | Adenocarcinoma |
| 15 | 49 | Male | Rectum | 2 | 1 | 3 | Adenocarcinoma |
| 16 | 39 | Female | Cecum | 1 | 1 | 2 | Non-adenocarcinoma |
| 17 | 35 | Male | Rectum | / | / | / | / |
| 18 | 71 | Male | Liver area | 1 | 2 | 4 | Non-adenocarcinoma |
| 19 | 56 | Male | Rectum | 2 | 1 | 3 | Adenocarcinoma |
| 20 | 47 | Male | Descending colon | 1 | 1 | 2 | Non-adenocarcinoma |
| 21 | 64 | Male | Rectum | 2 | 1 | 3 | Non-adenocarcinoma |
| 22 | 56 | Male | Ascending colon | 2 | 1 | 3 | Non-adenocarcinoma |
| 23 | 39 | Female | Rectum | 1 | 1 | 1 | Non-adenocarcinoma |

| | | | | | | | |
|---|---|---|---|---|---|---|---|
| 24 | 74 | Female | Rectum | 1 | 1 | 2 | Adenocarcinoma |
| 25 | 61 | Female | Rectum | 2 | 1 | 3 | Non-adenocarcinoma |
| 26 | / | Female | / | / | / | / | / |
| 27 | 56 | Female | Sigmoid colon | 2 | 1 | 3 | Non-adenocarcinoma |
| 28 | 50 | Female | Sigmoid colon | 1 | 1 | 2 | Adenocarcinoma |
| 29 | 68 | Female | Ascending colon | 2 | 1 | 3 | Adenocarcinoma |
| 30 | 56 | Female | Rectum | 2 | 1 | 3 | Non-adenocarcinoma |
| 31 | 70 | Female | Rectum | 1 | 1 | 1 | Adenocarcinoma |
| 32 | 67 | Male | Sigmoid colon | 1 | 1 | 2 | Adenocarcinoma |
| 33 | 71 | Female | Sigmoid colon | 2 | 1 | 3 | Non-adenocarcinoma |
| 34 | 51 | Male | Rectum | 1 | 1 | 2 | Adenocarcinoma |
| 35 | 56 | Male | Rectum | 2 | 1 | 3 | Non-adenocarcinoma |
| 36 | 55 | Female | Spleen area | 1 | 1 | 2 | Adenocarcinoma |
| 37 | 65 | Male | Rectum | 1 | 1 | 2 | Adenocarcinoma |
| 38 | 47 | Male | Sigmoid colon | 1 | 1 | 2 | Adenocarcinoma |
| 39 | 60 | Male | Sigmoid colon | 2 | 1 | 3 | Adenocarcinoma |
| 40 | 73 | Female | Transverse colon | 2 | 1 | 3 | Adenocarcinoma |
| 41 | 57 | Female | Sigmoid colon | 1 | 1 | 2 | Adenocarcinoma |
| 42 | 61 | Male | Sigmoid colon | 1 | 2 | 4 | Adenocarcinoma |
| 43 | 49 | Female | Rectum | 1 | 1 | 2 | Adenocarcinoma |
| 44 | 67 | Female | Rectum | 1 | 1 | 1 | Adenocarcinoma |
| 45 | 58 | Male | Sigmoid colon | 1 | 1 | 2 | Adenocarcinoma |
| 46 | 59 | Male | Spleen area | 2 | 1 | 3 | Adenocarcinoma |
| 47 | / | / | / | / | / | / | / |
| 48 | 72 | Male | Rectum | 1 | 1 | 2 | Adenocarcinoma |
| 49 | 55 | Male | Rectum | 1 | 1 | 2 | Non-adenocarcinoma |

| | | | | | | | |
|---|---|---|---|---|---|---|---|
| 50 | 54 | Male | Rectum | 1 | 1 | 2 | Non-adenocarcinoma |
| 51 | 74 | Female | Spleen area | 1 | 1 | 2 | Non-adenocarcinoma |

Note: Missing data: age ,2; sex, 2; tissue location, 2; T Stage, 4; M Stage, 4; N Stage, 4; TNM Stage, 4; Pathological type,4; histological_type, 4.

**Supplementary Table 2. Colorectal cancer metabolites identified from the paired-tissue metabolomics study**

| NO. | RT[a] | Mass | HMDB number | Compounds | Adduct type | ESI mode | Tendency (Case/Control) |
|---|---|---|---|---|---|---|---|
| Nucleotide | | | | | | | |
| 1 | 322.0385 | 0.97 | HMDB0000095 | Cytidine monophosphate | M-H | ESI- | ↑ |
| 2 | 346.0424 | 0.97 | HMDB0001397 | Guanosine monophosphate | M+H-H2O | ESI+ | ↑ |
| 3 | 352.0486 | 1.00 | HMDB0001202 | dCMP | M+FA-H | ESI- | ↑ |
| 4 | 345.0069 | 1.05 | HMDB0001554 | Xanthylic acid | / | ESI+/ESI- | ↑ |
| 5 | 346.0500 | 1.14 | HMDB0000045 | Adenosine monophosphate | M-H | ESI- | ↑ |
| 6 | 346.0483 | 1.88 | HMDB0003540 | 3'-AMP | / | ESI+/ESI- | ↑ |
| 7 | 374.0459 | 1.88 | HMDB0000058 | Cyclic AMP | / | ESI+/ESI- | ↑ |
| 8 | 376.0606 | 1.90 | HMDB0059612 | 7-Methylguanosine 5'-phosphate | / | ESI+/ESI- | ↑ |
| 9 | 434.0699 | 1.91 | HMDB0006880 | Acetyl adenylate | M+FA-H | ESI- | ↑ |
| 10 | 240.0907 | 1.97 | HMDB0002224 | 5-Methyldeoxycytidine | M-H | ESI- | ↑ |
| 11 | 376.0652 | 2.01 | HMDB0000905 | Deoxyadenosine monophosphate | M+FA-H | ESI- | ↑ |
| 12 | 346.0572 | 2.02 | HMDB0001044 | 2'-Deoxyguanosine 5'-monophosphate | / | ESI+/ESI- | ↑ |
| Amino acid | | | | | | | |
| 13 | 198.0153 | 0.70 | HMDB0006802 | O-Phospho-4-hydroxy-L-threonine | M+H-H2O | ESI+ | ↑ |
| 14 | 273.1085 | 2.09 | HMDB0000052 | Argininosuccinic acid | M+H-H2O | ESI+ | ↑ |
| 15 | 233.1298 | 2.54 | HMDB0000670 | Homo-L-arginine | M+FA-H | ESI- | ↓ |
| 16 | 187.0796 | 3.30 | HMDB0000929 | L-Tryptophan | M+H-H2O | ESI+ | ↑ |
| Carbohydrates | | | | | | | |
| 17 | 291.0708 | 0.78 | HMDB0000497 | 5,6-Dihydrouridine | M+FA-H | ESI- | ↑ |
| 18 | 403.0565 | 0.78 | HMDB0001124 | Trehalose 6-phosphate | / | ESI+/ESI- | ↑ |
| 19 | 210.9931 | 1.01 | HMDB0000618 | D-Ribulose 5-phosphate | M-H20-H | ESI- | ↑ |

| | | | | | | | |
|---|---|---|---|---|---|---|---|
| 20 | 181.0722 | 1.28 | HMDB0000122 | D-Glucose | M+H | ESI+ | ↑ |
| 21 | 210.9934 | 1.28 | HMDB0001548 | D-Ribose 5-phosphate | M-H20-H | ESI- | ↑ |
| 22 | 403.0716 | 1.82 | HMDB0006789 | Lactose 6-phosphate | M-H20-H | ESI- | ↑ |
| 23 | 210.9929 | 1.89 | HMDB0000868 | Xylulose 5-phosphate | M-H20-H | ESI- | ↑ |
| 24 | 404.0552 | 1.90 | HMDB0038413 | 2-Hydroxypentyl glucosinolate | M-H | ESI- | ↑ |
| 25 | 358.0505 | 1.94 | HMDB0000999 | Phosphoribosylformylglycineamidine | M+FA-H | ESI- | ↑ |
| 26 | 255.1035 | 1.95 | HMDB0000855 | Nicotinamide riboside | M+H | ESI+ | ↑ |
| 27 | 210.9933 | 2.01 | HMDB0001489 | Ribose 1-phosphate | M-H20-H | ESI- | ↑ |
| 28 | 360.0788 | 2.02 | HMDB0030934 | Triglochinin | M+H | ESI+ | ↑ |
| 29 | 657.2433 | 2.10 | HMDB0001081 | (N-acetylneuraminosyl(a2-6)lactosamine) | M+H-H2O | ESI+ | ↑ |
| Dipeptide | | | | | | | |
| 30 | 231.1684 | 0.97 | HMDB0028942 | Leucyl-Valine | M+H | ESI+ | ↑ |
| 31 | 203.1008 | 0.98 | HMDB0028688 | Alanyl-Hydroxyproline | M+H | ESI+ | ↑ |
| 32 | 221.0840 | 0.98 | HMDB0028848 | Glycyl-Phenylalanine | M-H | ESI- | ↓ |
| 33 | 211.1107 | 1.00 | HMDB0006695 | Prolylhydroxyproline | | ESI+ | ↑ |
| 34 | 286.1627 | 1.87 | HMDB0013328 | Pimelylcarnitine | M+H-H2O | ESI+ | ↑ |
| 35 | 239.1036 | 1.90 | HMDB0029105 | Tyrosyl-Glycine | M+H | ESI+ | ↑ |
| 36 | 447.1004 | 1.91 | HMDB0001117 | 4'-Phosphopantothenoylcysteine | M+FA-H | ESI- | ↑ |
| 37 | 205.0828 | 1.92 | HMDB0011667 | gamma-Glutamylglycine | M+H | ESI+ | ↑ |
| 38 | 173.0382 | 1.93 | HMDB0028684 | Alanyl-Cysteine | M-H20-H | ESI- | ↓ |
| 39 | 173.0497 | 1.94 | HMDB0028753 | Aspartyl-Glycine | M+H-H2O | ESI+ | ↓ |
| 40 | 213.1007 | 1.95 | HMDB0028885 | Histidinyl-Glycine | M+H | ESI+ | ↑ |
| 41 | 284.1355 | 1.95 | HMDB0028799 | Glutaminylhistidine | M+H | ESI+ | ↑ |
| 42 | 199.1017 | 1.99 | HMDB0029027 | Prolyl-Threonine | M+H-H2O | ESI+ | ↑ |
| 43 | 270.1091 | 2.05 | HMDB0028705 | Arginyl-Aspartic acid | M-H20-H | ESI- | ↑ |
| 44 | 217.1034 | 2.08 | HMDB0028778 | Cysteinyl-Isoleucine | M+H-H2O | ESI+ | ↓ |

| | | | | | | | |
|---|---|---|---|---|---|---|---|
| 45 | 234.1138 | 2.10 | HMDB0028741 | Asparaginyl-Threonine | M+H | ESI+ | ↑ |
| 46 | 270.1651 | 2.14 | HMDB0028710 | Arginyl-Hydroxyproline | M+H-H2O | ESI+ | ↑ |
| 47 | 286.1076 | 2.15 | HMDB0029072 | Threoninyl-Tryptophan | M-H20-H | ESI- | ↑ |
| 48 | 209.1019 | 2.19 | HMDB0028689 | Alanyl-Histidine | M+H-H2O | ESI+ | ↑ |
| 49 | 233.1397 | 2.21 | HMDB0028939 | Leucyl-Threonine | M+H | ESI+ | ↑ |
| 50 | 213.0832 | 2.23 | HMDB0028766 | Aspartyl-Valine | M-H20-H | ESI- | ↑ |
| 51 | 243.1246 | 2.26 | HMDB0011177 | Phenylalanylproline | / | ESI+/ESI- | ↑ |
| 52 | 233.0643 | 2.37 | HMDB0028774 | Cysteinyl-Glutamate | M+H-H2O | ESI+ | ↓ |
| 53 | 299.1081 | 2.47 | HMDB0028742 | Asparaginyl-Tryptophan | M-H20-H | ESI- | ↓ |
| 54 | 284.169 | 2.51 | HMDB0028890 | Histidinyl-Lysine | M+H | ESI+ | ↑ |
| 55 | 291.0946 | 2.54 | HMDB0011741 | gamma-Glutamyltyrosine | M-H20-H | ESI- | ↓ |
| 56 | 312.1927 | 2.56 | HMDB0028711 | Arginyl-Histidine | M+H | ESI+ | ↑ |
| 57 | 269.0981 | 2.57 | HMDB0028755 | Aspartyl-Histidine | M-H | ESI- | ↓ |
| 58 | 247.1105 | 2.58 | HMDB0028986 | Methionyl-Valine | M-H | ESI- | ↓ |
| 59 | 257.1363 | 2.58 | HMDB0029063 | Threoninyl-Histidine | M+H | ESI+ | ↓ |
| 60 | 298.1548 | 2.62 | HMDB0028918 | Isoleucyl-Tryptophan | M-H20-H | ESI- | ↑ |
| 61 | 267.1073 | 2.70 | HMDB0028995 | Phenylalanyl-Glycine | M+FA-H | ESI- | ↓ |
| 62 | 242.1106 | 2.74 | HMDB0028877 | Hydroxyprolyl-Gamma-glutamate | M+H-H2O | ESI+ | ↓ |
| 63 | 247.0987 | 2.74 | HMDB0028726 | Asparaginyl-Asparagine | M+H | ESI+ | ↓ |
| 64 | 332.1255 | 2.84 | HMDB0028830 | Glutamyltryptophan | M-H | ESI- | ↑ |
| 65 | 299.1348 | 2.90 | HMDB0028898 | Histidinyl-Valine | M+FA-H | ESI- | ↓ |
| 66 | 346.1473 | 2.94 | HMDB0029028 | Prolyl-Tryptophan | M+FA-H | ESI- | ↑ |
| 67 | 366.1659 | 2.95 | HMDB0028716 | Arginyl-Phenylalanine | M+FA-H | ESI- | ↑ |
| 68 | 277.1514 | 2.97 | HMDB0013243 | Leucyl-phenylalanine | M-H | ESI- | ↓ |
| 69 | 259.1429 | 3.02 | HMDB0011171 | gamma-Glutamylleucine | M-H | ESI- | ↓ |
| 70 | 319.1678 | 3.02 | HMDB0028965 | Lysyl-Gamma-glutamate | M+FA-H | ESI- | ↓ |

| | | | | | | | |
|---|---|---|---|---|---|---|---|
| 71 | 243.1332 | 3.13 | HMDB0028928 | Leucyl-Glutamate | M+H-H2O | ESI+ | ↑ |
| 72 | 225.1071 | 3.28 | HMDB0028878 | Histidinyl-Alanine | / | ESI+/ESI- | ↓ |
| 73 | 243.1162 | 3.28 | HMDB0011179 | Prolylphenylalanine | M-H20-H | ESI- | ↓ |
| 74 | 295.1187 | 3.29 | HMDB0028980 | Methionyl-Phenylalanine | M-H | ESI- | ↓ |
| 75 | 249.0978 | 3.30 | HMDB0029159 | gamma-Glutamylthreonine | M+H | ESI+ | ↓ |
| 76 | 261.1268 | 3.49 | HMDB0028996 | Phenylalanyl-Hydroxyproline | M+H-H2O | ESI+ | ↑ |
| Lipids | | | | | | | |
| 77 | 469.2104 | 2.28 | HMDB0002421 | 7-Sulfocholic acid | M-H20-H | ESI- | ↑ |
| 78 | 441.2492 | 2.93 | HMDB0002504 | 3-Sulfodeoxycholic acid | M+H-H2O | ESI+ | ↑ |
| 79 | 415.2179 | 3.18 | HMDB0002886 | 6-Keto-prostaglandin F1a | M+FA-H | ESI- | ↓ |
| 80 | 363.2151 | 3.29 | HMDB0001337 | Leukotriene A4 | M+FA-H | ESI- | ↓ |
| 81 | 467.2580 | 3.43 | HMDB0010321 | 3,17-Androstanediol glucuronide | M-H | ESI- | ↑ |
| 82 | 423.2035 | 3.47 | HMDB0000418 | 18-Hydroxycortisol | M+FA-H | ESI- | ↓ |
| 83 | 387.2142 | 3.48 | HMDB0002277 | 2,3-Dinor-6-keto-prostaglandin F1 a | M+FA-H | ESI- | ↓ |
| 84 | 397.2233 | 3.49 | HMDB0001335 | Prostaglandin I2 | M+FA-H | ESI- | ↑ |
| 85 | 481.2605 | 3.57 | HMDB0010351 | 11-beta-Hydroxyandrosterone-3-glucuronide | M-H | ESI- | ↑ |
| 86 | 355.2401 | 3.64 | HMDB0004239 | 13,14-Dihydro PGF2a | M-H | ESI- | ↑ |
| 87 | 379.2350 | 3.70 | HMDB0002082 | Bisnorcholic acid | M-H | ESI- | ↑ |
| 88 | 319.2149 | 3.72 | HMDB0002265 | 14,15-DiHETrE | M-H20-H | ESI- | ↓ |
| 89 | 325.2191 | 3.78 | HMDB0010213 | 17 HDoHE | M-H20-H | ESI- | ↓ |
| 90 | 381.2458 | 3.80 | HMDB0001085 | Leukotriene B4 | M+FA-H | ESI- | ↑ |
| 91 | 367.2180 | 3.86 | HMDB0000385 | 12a-Hydroxy-3-oxocholadienic acid | M-H20-H | ESI- | ↑ |
| 92 | 381.2487 | 3.92 | HMDB0012518 | 11'-Carboxy-gamma-tocotrienol | M-H20-H | ESI- | ↑ |
| 93 | 313.2299 | 3.95 | HMDB0000315 | 16-A-Hydroxypregnenolone | M-H20-H | ESI- | ↓ |
| 94 | 480.2997 | 4.00 | HMDB0000874 | Tauroursodeoxycholic acid | M-H20-H | ESI- | ↑ |
| 95 | 313.2299 | 4.02 | HMDB0000363 | 17a-Hydroxypregnenolone | M-H20-H | ESI- | ↓ |

| | | | | | | | |
|---|---|---|---|---|---|---|---|
| 96 | 409.2400 | 4.09 | HMDB0003259 | Dihydrocortisol | M+FA-H | ESI- | ↑ |
| 97 | 201.1136 | 4.10 | HMDB0000792 | Sebacic acid | M-H | ESI- | ↓ |
| 98 | 703.3375 | 4.12 | HMDB0036338 | 25-Acetyl-6,7-didehydrofevicordin F 3-glucoside | M-H | ESI- | ↑ |
| 99 | 411.2367 | 4.14 | HMDB0000949 | Tetrahydrocortisol | M+FA-H | ESI- | ↑ |
| 100 | 315.2439 | 4.26 | HMDB0000253 | Pregnenolone | M-H | ESI- | ↓ |
| 101 | 315.2432 | 4.39 | HMDB0011532 | MG(0:0/15:0/0:0) | M-H | ESI- | ↓ |
| 102 | 373.2056 | 4.40 | HMDB0007003 | CPA(16:0/0:0) | M-H20-H | ESI- | ↓ |
| 103 | 438.2101 | 4.42 | HMDB0012639 | 20-Hydroxy-leukotriene E4 | M+H-H2O | ESI+ | ↑ |
| 104 | 353.2269 | 4.54 | HMDB0001220 | Prostaglandin E2 | M+H | ESI+ | ↓ |
| 105 | 295.2176 | 4.69 | HMDB0004667 | 13S-hydroxyoctadecadienoic acid | M-H | ESI- | ↑ |
| 106 | 613.4435 | 4.69 | HMDB0071165 | TG(10:0/13:0/8:0) | M+FA-H | ESI- | ↑ |
| 107 | 830.6059 | 4.74 | HMDB0000593 | PC(18:1(9Z)/18:1(9Z)) | M+FA-H | ESI- | ↑ |
| 108 | 219.1323 | 4.76 | HMDB0036389 | 3-Phenylpropyl isovalerate | M-H | ESI- | ↑ |
| 109 | 345.2443 | 4.93 | HMDB0011531 | MG(0:0/14:1(9Z)/0:0) | M+FA-H | ESI- | ↑ |
| 110 | 345.2381 | 5.05 | HMDB0011562 | MG(14:1(9Z)/0:0/0:0) | M+FA-H | ESI- | ↑ |
| 111 | 293.2031 | 5.18 | HMDB0004668 | 13-OxoODE | M-H | ESI- | ↑ |
| 112 | 800.5935 | 5.35 | HMDB0007952 | PC(15:0/22:1(13Z)) | M-H | ESI- | ↑ |
| 113 | 267.2247 | 5.42 | HMDB0030997 | Cyclohexaneundecanoic acid | M-H | ESI- | ↑ |
| 114 | 271.2193 | 5.43 | HMDB0010734 | (R)-3-Hydroxy-hexadecanoic acid | M-H | ESI- | ↓ |
| 115 | 323.2506 | 5.65 | HMDB0029797 | (Z)-15-Oxo-11-eicosenoic acid | M-H | ESI- | ↑ |
| 116 | 305.2160 | 5.73 | HMDB0013156 | 16-alpha-Hydroxyandrosterone | M-H | ESI- | ↑ |
| 117 | 317.2058 | 5.75 | HMDB0010202 | 12-HEPE | M-H | ESI- | ↑ |
| 118 | 299.2485 | 5.88 | HMDB0061661 | 9-hydroxyoctadecanoic acid | M-H | ESI- | ↓ |
| 119 | 496.3347 | 6.02 | HMDB0010382 | LysoPC(16:0) | M+H | ESI+ | ↑ |
| 120 | 217.1328 | 6.63 | HMDB0000888 | Undecanedioic acid | M+H | ESI+ | ↓ |
| 121 | 546.3512 | 6.93 | HMDB0010393 | LysoPC(20:3(5Z,8Z,11Z)) | M+H | ESI+ | ↑ |

| | | | | | | | |
|---|---|---|---|---|---|---|---|
| 122 | 301.2106 | 7.00 | HMDB0001999 | Eicosapentaenoic acid | M-H | ESI- | ↑ |
| 123 | 355.2823 | 7.92 | HMDB0011538 | MG(0:0/18:2(9Z,12Z)/0:0) | M+H | ESI+ | ↓ |
| 124 | 395.2756 | 7.93 | HMDB0012530 | 11-Hydroxyeicosatetraenoate glyceryl ester | M+H | ESI+ | ↓ |
| 125 | 414.3224 | 7.93 | HMDB0013333 | 3-Hydroxy-9-hexadecenoylcarnitine | M+H | ESI+ | ↓ |
| 126 | 283.2352 | 8.03 | HMDB0011530 | MG(0:0/14:0/0:0) | M-H20-H | ESI- | ↑ |
| 127 | 395.2216 | 8.04 | HMDB0002664 | Prostaglandin E3 | M+FA-H | ESI- | ↑ |
| 128 | 253.2086 | 8.23 | HMDB0003229 | Palmitoleic acid | M-H | ESI- | ↑ |
| 129 | 529.4244 | 8.23 | HMDB0092933 | DG(8:0/18:0/0:0) | M+FA-H | ESI- | ↑ |
| 130 | 253.2095 | 8.40 | HMDB0031053 | (E)-6-Hexadecenoic acid | M-H | ESI- | ↑ |
| 131 | 205.1884 | 8.53 | HMDB0040377 | 3-Pentadecenal | M-H20-H | ESI- | ↑ |
| 132 | 303.2213 | 8.53 | HMDB0001043 | Arachidonic acid | / | ESI+/ESI- | ↑ |
| 133 | 361.1857 | 8.53 | HMDB0000319 | 18-Hydroxycorticosterone | M-H | ESI- | ↑ |
| 134 | 629.4581 | 8.54 | HMDB0000674 | PA(16:0/16:0) | M-H20-H | ESI- | ↑ |
| 135 | 361.1852 | 8.61 | HMDB0000063 | Cortisol | M-H | ESI- | ↑ |
| 136 | 329.2418 | 8.90 | HMDB0004708 | 9,12,13-TriHOME | M-H | ESI- | ↑ |
| 137 | 559.4719 | 8.91 | HMDB0007074 | DG(15:0/18:2(9Z,12Z)/0:0) | M-H20-H | ESI- | ↑ |
| 138 | 267.2256 | 9.46 | HMDB0031046 | 9E-Heptadecenoic acid | M-H | ESI- | ↑ |
| 139 | 329.2419 | 9.48 | HMDB0004710 | 9,10,13-TriHOME | M-H | ESI- | ↑ |
| 140 | 305.2399 | 9.54 | HMDB0010378 | 5,8,11-Eicosatrienoic acid | M-H | ESI- | ↑ |
| 141 | 529.4076 | 9.54 | HMDB0006816 | 3-Hexaprenyl-4-hydroxybenzoic acid | M+H-H2O | ESI+ | ↓ |
| 142 | 633.4935 | 9.55 | HMDB0007025 | DG(14:0/20:4(5Z,8Z,11Z,14Z)/0:0) | M+FA-H | ESI- | ↑ |
| 143 | 521.3729 | 9.89 | HMDB0114757 | LysoPA(24:1(15Z)/0:0) | M+H | ESI+ | ↓ |
| 144 | 331.2562 | 10.05 | HMDB0002226 | Adrenic acid | M-H | ESI- | ↑ |
| 145 | 509.3840 | 10.39 | HMDB0035150 | 2'-Apo-beta-carotenal | M+H | ESI+ | ↓ |
| Others | | | | | | | |
| 146 | 137.0431 | 1.02 | HMDB0000157 | Hypoxanthine | M+H | ESI+ | ↑ |

| | | | | | | | |
|---|---|---|---|---|---|---|---|
| 147 | 146.0971 | 1.02 | HMDB0003464 | 4-Guanidinobutanoic acid | M+H | ESI+ | ↑ |
| 148 | 249.0534 | 1.06 | HMDB0001555 | Pyridoxamine 5'-phosphate | M+H | ESI+ | ↑ |
| 149 | 256.1012 | 1.14 | HMDB0002275 | 7,8-Dihydroneopterin | M+H | ESI+ | ↑ |
| 150 | 136.0602 | 1.86 | HMDB0000034 | Adenine | M+H | ESI+ | ↑ |
| 151 | 523.0658 | 1.87 | HMDB0029212 | Quercetin 3-O-glucuronide | M+FA-H | ESI- | ↑ |
| 152 | 164.0564 | 1.90 | HMDB0006037 | 8-Hydroxy-7-methylguanine | M+H-H2O | ESI+ | ↑ |
| 153 | 449.1183 | 1.92 | HMDB0029209 | naringenin-7-O-glucuronide | M+H | ESI+ | ↑ |
| 154 | 271.0886 | 1.97 | HMDB0000245 | Porphobilinogen | M+FA-H | ESI- | ↑ |
| 155 | 483.3668 | 10.31 | HMDB0032111 | Adlupone | M+H | ESI+ | ↓ |
| 156 | 180.0894 | 2.10 | HMDB0011690 | 7-Aminomethyl-7-carbaguanine | M+H | ESI+ | ↑ |
| 157 | 148.0614 | 2.12 | HMDB0001566 | 3-Methylguanine | M+H-H2O | ESI+ | ↑ |
| 158 | 304.0941 | 2.14 | HMDB0029839 | Ascorbigen | M-H | ESI- | ↑ |
| 159 | 241.0640 | 2.31 | HMDB0001857 | 1,3-Dimethyluric acid | M+FA-H | ESI- | ↑ |
| 160 | 263.0971 | 2.31 | HMDB0000235 | Thiamine | M-H | ESI- | ↑ |
| 161 | 414.1922 | 2.35 | HMDB0010341 | Dextrorphan O-glucuronide | M-H20-H | ESI- | ↑ |
| 162 | 174.0921 | 2.52 | HMDB0004225 | 2-Oxoarginine | M+H | ESI+ | ↑ |
| 163 | 127.0697 | 2.75 | HMDB0003701 | Dimethylbenzimidazole | M-H20-H | ESI- | ↓ |
| 164 | 239.0907 | 2.76 | HMDB0031517 | (R)-3-Hydroxy-5-phenylpentanoic acid | M+FA-H | ESI- | ↓ |
| 165 | 385.1649 | 2.90 | HMDB0001091 | 3 Hydroxyquinine | M+FA-H | ESI- | ↑ |
| 166 | 217.1000 | 2.91 | HMDB0000350 | 3-Hydroxysebacic acid | M-H | ESI- | ↓ |
| 167 | 337.1363 | 3.17 | HMDB0011687 | Phenylbutyrylglutamine | M+FA-H | ESI- | ↑ |
| 168 | 233.1314 | 3.20 | HMDB0010725 | (R)-3-Hydroxydecanoic acid | M+FA-H | ESI- | ↓ |
| 169 | 167.1014 | 3.60 | HMDB0010724 | 3-Oxodecanoic acid | M-H20-H | ESI- | ↑ |
| 170 | 233.1454 | 3.63 | HMDB0002203 | 3-Hydroxycapric acid | M+FA-H | ESI- | ↑ |
| 171 | 195.1314 | 4.69 | HMDB0010727 | 3-Oxododecanoic acid | M-H20-H | ESI- | ↑ |
| 172 | 171.0940 | 4.86 | HMDB0004095 | 5-Methoxytryptamine | M-H20-H | ESI- | ↑ |

| | | | | | | | |
|---|---|---|---|---|---|---|---|
| 173 | 259.2356 | 8.53 | HMDB0033609 | 2- Pentadecylfuran | M-H20-H | ESI- | ↑ |

[a]RT, retention time

**Supplementary Table 3. Pathway analysis of 373 positive and negative ion by Mummichog**

| Pathway | Overlap_size | Pathway_size | *P*-value (raw) | *P*-value |
|---|---|---|---|---|
| **Negative model** | | | | |
| Linoleate metabolism | 17 | 21 | 0.00012 | 0.005693 |
| Leukotriene metabolism | 31 | 50 | 0.000939 | 0.00573 |
| Drug metabolism - cytochrome P450 | 27 | 48 | 0.013012 | 0.006287 |
| Fatty acid activation | 12 | 18 | 0.017944 | 0.006951 |
| Prostaglandin formation from arachidonate | 33 | 64 | 0.031624 | 0.006997 |
| De novo fatty acid biosynthesis | 11 | 17 | 0.031337 | 0.007887 |
| Prostaglandin formation from dihomo gama-linoleic acid | 5 | 6 | 0.038748 | 0.01118 |
| Limonene and pinene degradation | 5 | 6 | 0.038748 | 0.01118 |
| Biopterin metabolism | 9 | 15 | 0.088274 | 0.012531 |
| Vitamin A (retinol) metabolism | 18 | 36 | 0.131433 | 0.013277 |
| Omega-3 fatty acid metabolism | 5 | 7 | 0.091712 | 0.017255 |
| Vitamin B5 - CoA biosynthesis from pantothenate | 6 | 10 | 0.158468 | 0.023387 |
| N-Glycan Degradation | 4 | 6 | 0.172934 | 0.034215 |
| Dynorphin metabolism | 4 | 6 | 0.172934 | 0.034215 |
| CoA Catabolism | 4 | 6 | 0.172934 | 0.034215 |
| Pentose phosphate pathway | 13 | 29 | 0.343748 | 0.041055 |
| **Positive model** | | | | |
| N-Glycan Degradation | 6 | 8 | 0.017992 | 0.00075 |
| Drug metabolism - cytochrome P450 | 22 | 51 | 0.07712 | 0.000995 |
| Sialic acid metabolism | 13 | 28 | 0.092295 | 0.001332 |
| Heparan sulfate degradation | 4 | 5 | 0.042719 | 0.001607 |
| Vitamin B5 - CoA biosynthesis from pantothenate | 5 | 8 | 0.082653 | 0.002154 |

| | | | | |
|---|---|---|---|---|
| Chondroitin sulfate degradation | 3 | 3 | 0.035349 | 0.002221 |
| Glycosphingolipid metabolism | 11 | 25 | 0.162942 | 0.00271 |
| Prostaglandin formation from dihomo gama-linoleic acid | 4 | 6 | 0.09513 | 0.003208 |
| Leukotriene metabolism | 19 | 50 | 0.260187 | 0.004551 |
| Omega-6 fatty acid metabolism | 3 | 4 | 0.106692 | 0.005744 |
| Vitamin H (biotin) metabolism | 3 | 4 | 0.106692 | 0.005744 |
| Tryptophan metabolism | 22 | 60 | 0.304843 | 0.005954 |
| Biopterin metabolism | 6 | 14 | 0.296041 | 0.011921 |
| Prostaglandin formation from arachidonate | 22 | 63 | 0.408527 | 0.012283 |
| Omega-3 fatty acid metabolism | 4 | 8 | 0.248651 | 0.012868 |
| Keratan sulfate degradation | 4 | 8 | 0.248651 | 0.012868 |
| Histidine metabolism | 9 | 24 | 0.385836 | 0.016438 |
| Butanoate metabolism | 7 | 18 | 0.374535 | 0.018158 |
| Glycerophospholipid metabolism | 13 | 37 | 0.444616 | 0.019751 |
| N-Glycan biosynthesis | 5 | 12 | 0.354775 | 0.021228 |
| Aminosugars metabolism | 10 | 28 | 0.443216 | 0.022564 |
| Aspartate and asparagine metabolism | 17 | 50 | 0.485278 | 0.022807 |
| De novo fatty acid biosynthesis | 7 | 19 | 0.439384 | 0.027748 |
| CoA Catabolism | 3 | 6 | 0.310175 | 0.029356 |
| Glycosphingolipid biosynthesis - globoseries | 3 | 6 | 0.310175 | 0.029356 |
| Glycolysis and Gluconeogenesis | 7 | 20 | 0.503064 | 0.041215 |
| Vitamin B1 (thiamin) metabolism | 4 | 10 | 0.427937 | 0.041983 |
| Glycosphingolipid biosynthesis - ganglioseries | 4 | 10 | 0.427937 | 0.041983 |

**Supplementary Table 4. Fifty identifiable metabolites differential abundance between early (I, II) and late-stage (III, IV) tumors.**

| NO. | RT[a] | Mass | HMDB number | Compounds | Adduct type | ESI mode |
|---|---|---|---|---|---|---|
| 1 | 294.0953 | 2.32 | HMDB0028743 | Asparaginyl-Tyrosine | ESI- | M-H |
| 2 | 321.1433 | 2.48 | HMDB0033105 | N2-Galacturonyl-L-lysine | ESI- | M-H |
| 3 | 277.1203 | 2.48 | HMDB0011176 | L-phenylalanyl-L-hydroxyproline | ESI- | M-H |
| 4 | 233.1298 | 2.54 | HMDB0000670 | Homo-L-arginine | ESI- | M+FA-H |
| 5 | 245.1304 | 2.71 | HMDB0029008 | Phenylalanyl-Valine | ESI- | M-H-H2O |
| 6 | 291.1347 | 2.73 | HMDB0028887 | Histidinyl-Histidine | ESI- | M-H |
| 7 | 247.1161 | 2.74 | HMDB0028986 | Methionyl-Valine | ESI- | M-H |
| 8 | 339.1442 | 2.77 | HMDB0028919 | Isoleucyl-Tyrosine | ESI- | M+FA-H |
| 9 | 261.1261 | 2.81 | HMDB0028913 | Isoleucyl-Methionine | ESI- | M-H |
| 10 | 217.1 | 2.91 | HMDB0028694 | Alanyl-Phenylalanine | ESI- | M-H-H2O |
| 11 | 237.1055 | 2.95 | HMDB0028895 | Histidinyl-Threonine | ESI- | M-H-H2O |
| 12 | 187.0882 | 3.1 | HMDB0059783 | 3-Methylsubericacid | ESI- | M-H |
| 13 | 345.216 | 3.2 | HMDB0006219 | 13-cis-retinoic acid,Isotretinoin | ESI- | M+FA-H |
| 14 | 363.2151 | 3.29 | HMDB0010209 | 15-HEPE | ESI- | M+FA-H |
| 15 | 285.1422 | 3.35 | HMDB0000313 | 16b-Hydroxyestrone | ESI- | M-H |
| 16 | 169.1154 | 3.36 | HMDB0012183 | 8-Methylnonenoate | ESI- | M-H |
| 17 | 235.1071 | 3.36 | HMDB0028988 | Phenylalanyl-Alanine | ESI- | M-H |
| 18 | 273.1254 | 3.37 | HMDB0011687 | Phenylbutyrylglutamine | ESI- | M-H-H2O |
| 19 | 311.1819 | 3.39 | HMDB0028703 | Arginyl-Arginine | ESI- | M-H-H2O |
| 20 | 373.2189 | 3.39 | HMDB0007003 | CPA(16:0/0:0) | ESI- | M-H-H2O |
| 21 | 285.1807 | 3.55 | HMDB0000352 | 16alpha-Hydroxydehydroepiandrosterone | ESI- | M-H-H2O |
| 22 | 285.1713 | 3.61 | HMDB0000309 | 3a,16b-Dihydroxyandrostenone | ESI- | M-H-H2O |
| 23 | 287.2124 | 3.81 | HMDB0000077 | Dehydroepiandrosterone | ESI- | M-H |

| | | | | | | |
|---|---|---|---|---|---|---|
| 24 | 287.2136 | 3.91 | HMDB0000899 | 5alpha-androstane-3,17-dione | ESI- | M-H |
| 25 | 480.2997 | 4 | HMDB0011130 | PE(18:0/0:0) | ESI- | M-H |
| 26 | 241.1698 | 4.15 | HMDB0010730 | 3-oxo-tetradecanoic acid | ESI- | M-H |
| 27 | 269.2046 | 4.24 | HMDB0072845 | MG(13:0/0:0/0:0) | ESI- | M-H-H2O |
| 28 | 309.2006 | 4.36 | HMDB0013623 | 9-oxo-12,13-epoxy-10-octadecenoic acid | ESI- | M-H |
| 29 | 269.2047 | 4.36 | HMDB0072861 | MG(0:0/13:0/0:0) | ESI- | M-H-H2O |
| 30 | 554.3408 | 4.58 | HMDB0011511 | PE(20:0/0:0) | ESI- | M+FA-H |
| 31 | 494.3136 | 4.58 | HMDB0006898 | Chenodeoxyglycocholic acid | ESI- | M+FA-H |
| 32 | 544.3172 | 4.58 | HMDB0000951 | Taurochenodeoxycholic acid | ESI- | M+FA-H |
| 33 | 494.3111 | 4.72 | HMDB0000708 | Glycoursodeoxycholic acid | ESI- | M+FA-H |
| 34 | 552.2899 | 4.75 | HMDB0011499 | LysoPE(0:0/24:6(6Z,9Z,12Z,15Z,18Z,21Z)) | ESI- | M-H |
| 35 | 297.2309 | 5.21 | HMDB0010736 | 3-keto stearic acid | ESI- | M-H |
| 36 | 265.1349 | 5.53 | HMDB0000372 | 16-Oxoestrone | ESI- | M-H-H2O |
| 37 | 311.154 | 6.41 | HMDB0061027 | 2-hydroxyethinylestradiol | ESI- | M-H |
| 38 | 297.2348 | 6.7 | HMDB0059633 | (9S,10S)-9,10-dihydroxyoctadecanoic acid | ESI- | M-H-H2O |
| 39 | 255.1517 | 3.31 | HMDB0028898 | Histidinyl-Valine | ESI+ | M+H |
| 40 | 313.2359 | 4.54 | HMDB0003871 | 13S-HpODE | ESI+ | M+H |
| 41 | 480.3422 | 6.87 | HMDB0010407 | PC(P-16:0/0:0) | ESI+ | M+H |
| 42 | 492.3383 | 6.87 | HMDB0011481 | LysoPE(0:0/20:0) | ESI+ | M+H-H2O |
| 43 | 510.3366 | 6.88 | HMDB0011511 | PE(20:0/0:0) | ESI+ | M+H |
| 44 | 518.3203 | 7.02 | HMDB0010387 | PC(18:3(6Z,9Z,12Z)/0:0) | ESI+ | M+H |
| 45 | 466.3273 | 7.02 | HMDB0000138 | Glycocholic acid | ESI+ | M+H |
| 46 | 478.3234 | 7.02 | HMDB0010382 | PC(16:0/0:0) | ESI+ | M+H-H2O |
| 47 | 482.3418 | 7.6 | HMDB0011129 | LysoPE(0:0/18:0) | ESI+ | M+H |
| 48 | 508.3546 | 7.82 | HMDB0011482 | LysoPE(0:0/20:1(11Z)) | ESI+ | M+H |
| 49 | 538.386 | 8.02 | HMDB0011490 | LysoPE(0:0/22:0) | ESI+ | M+H |

| | | | | | | |
|---|---|---|---|---|---|---|
| 50 | 385.2907 | 8.58 | HMDB0011557 | MG(0:0/22:6(4Z,7Z,10Z,13Z,16Z,19Z)/0:0) | ESI+ | M+H-H2O |

[a]RT, retention time

**Supplementary Table 5. Pathway analysis of 94 positive and negative ion by Mummichog**

| Pathway | Overlap_size | Pathway_size | *P*-value (raw) | *P*-value |
|---|---|---|---|---|
| **Negative model** | | | | |
| Linoleate metabolism | 7 | 21 | 0.073057 | 0.001555 |
| Androgen and estrogen biosynthesis and metabolism | 17 | 70 | 0.125102 | 0.001723 |
| Vitamin A (retinol) metabolism | 9 | 36 | 0.199974 | 0.003174 |
| Omega-3 fatty acid metabolism | 3 | 7 | 0.12024 | 0.003895 |
| De novo fatty acid biosynthesis | 5 | 17 | 0.186812 | 0.003948 |
| C21-steroid hormone biosynthesis and metabolism | 18 | 87 | 0.32218 | 0.004827 |
| Drug metabolism - cytochrome P450 | 10 | 48 | 0.382566 | 0.007938 |
| Urea cycle/amino group metabolism | 9 | 46 | 0.474658 | 0.012985 |
| Limonene and pinene degradation | 2 | 6 | 0.303833 | 0.019796 |
| Tryptophan metabolism | 11 | 68 | 0.729774 | 0.041297 |
| **Positive model** | | | | |
| Carnitine shuttle | 5 | 25 | 0.003321 | 0.001247 |
| Porphyrin metabolism | 2 | 22 | 0.24149 | 0.010312 |
| Bile acid biosynthesis | 3 | 54 | 0.409312 | 0.013294 |
| C21-steroid hormone biosynthesis and metabolism | 4 | 83 | 0.479562 | 0.014328 |
| Androgen and estrogen biosynthesis and metabolism | 3 | 66 | 0.544338 | 0.021534 |
| Vitamin E metabolism | 2 | 36 | 0.461136 | 0.022977 |
| Aspartate and asparagine metabolism | 2 | 50 | 0.641177 | 0.040161 |
| Leukotriene metabolism | 2 | 50 | 0.641177 | 0.040161 |

**Supplementary Table 6. Twenty-six identifiable metabolites differential abundance between adenocarcinoma and non-adenocarcinoma tumors.**

| NO. | RT[a] | Mass | HMDB number | Compounds | Adduct type | ESI mode |
|---|---|---|---|---|---|---|
| 1 | 298.0742 | 0.97 | HMDB0000845 | Neopterin | ESI- | M+FA-H |
| 2 | 213.0178 | 2.55 | HMDB0001351 | Deoxyribose 1-phosphate | ESI- | M-H |
| 3 | 347.2302 | 3.36 | HMDB0000949 | Tetrahydrocortisol | ESI- | M-H20-H |
| 4 | 255.1085 | 3.63 | HMDB0028795 | Glutaminylglutamine | ESI- | M-H20-H |
| 5 | 469.2467 | 3.94 | HMDB0002421 | 7-Sulfocholic acid | ESI- | M-H20-H |
| 6 | 301.2234 | 4.12 | HMDB0002190 | 5,6-Epoxy-8,11,14-eicosatrienoic acid | ESI- | M-H20-H |
| 7 | 353.2024 | 4.22 | HMDB0012869 | 9'-Carboxy-gamma-tocotrienol | ESI- | M-H20-H |
| 8 | 369.1847 | 4.34 | HMDB0002759 | Androsterone sulfate | ESI- | M-H |
| 9 | 476.2744 | 4.52 | HMDB0011477 | LysoPE(0:0/18:2(9Z,12Z)) | ESI- | M-H |
| 10 | 516.3209 | 5.3 | HMDB0010387 | LysoPC(18:3(6Z,9Z,12Z)) | ESI- | M-H |
| 11 | 542.3401 | 5.58 | HMDB0010395 | LysoPC(20:4(5Z,8Z,11Z,14Z)) | ESI- | M-H |
| 12 | 624.3024 | 5.58 | HMDB0001198 | Leukotriene C4 | ESI- | M-H |
| 13 | 323.1734 | 5.89 | HMDB0013243 | Leucyl-phenylalanine | ESI- | M+FA-H |
| 14 | 544.3543 | 7.1 | HMDB0010393 | LysoPC(20:3(5Z,8Z,11Z)) | ESI- | M-H |
| 15 | 554.3842 | 7.1 | HMDB0011149 | LysoPC(O-18:0) | ESI- | M+FA-H |
| 16 | 271.2208 | 7.57 | HMDB0010734 | (R)-3-Hydroxy-hexadecanoic acid | ESI- | M-H |
| 17 | 227.1938 | 7.77 | HMDB0002221 | 2,6,10-Trimethylundecanoic acid | ESI- | M-H |
| 18 | 241.2093 | 9.12 | HMDB0032250 | Dodecyl propionate | ESI- | M-H |
| 19 | 255.2242 | 10.1 | HMDB0031068 | Isopalmitic acid | ESI- | M-H |
| 20 | 345.2568 | 10.31 | HMDB0010737 | (R)-3-Hydroxy-Octadecanoic acid | ESI- | M+FA-H |
| 21 | 567.3612 | 10.33 | HMDB0114756 | LysoPA(24:0/0:0) | ESI- | M+FA-H |
| 22 | 511.3629 | 10.34 | HMDB0002972 | Vitamin K1 2,3-epoxide | ESI- | M+FA-H |
| 23 | 316.1186 | 2.3 | HMDB0028830 | Glutamyltryptophan | ESI+ | M+H-H2O |

| | | | | | | |
|---|---|---|---|---|---|---|
| 24 | 370.2865 | 4.56 | HMDB0013329 | trans-2-Tetradecenoylcarnitine | ESI+ | M+H |
| 25 | 263.2297 | 6.45 | HMDB0000673 | Linoleic acid | ESI+ | M+H-H2O |
| 26 | 285.1855 | 6.52 | HMDB0003955 | 19-Hydroxyandrost-4-ene-3,17-dione | ESI+ | M+H-H2O |

**Supplementary Table 7. Pathway analysis of 43 positive and negative ion by Mummichog**

| Pathway | Overlap_size | Pathway_size | *P*-value (raw) | *P*-value |
|---|---|---|---|---|
| Fatty acid activation | 4 | 18 | 0.092743 | 2.18E-05 |
| Fatty Acid Metabolism | 3 | 12 | 0.106204 | 5.09E-05 |
| De novo fatty acid biosynthesis | 3 | 17 | 0.23038 | 0.000374 |
| Carnitine shuttle | 3 | 21 | 0.342279 | 0.001478 |
| Vitamin E metabolism | 4 | 34 | 0.434647 | 0.002318 |
| Vitamin A (retinol) metabolism | 4 | 36 | 0.480125 | 0.003702 |
| Prostaglandin formation from arachidonate | 6 | 64 | 0.615755 | 0.008622 |
| Biopterin metabolism | 2 | 15 | 0.443342 | 0.011149 |
| Drug metabolism - cytochrome P450 | 4 | 48 | 0.712443 | 0.031625 |
| Leukotriene metabolism | 4 | 50 | 0.742804 | 0.041045 |

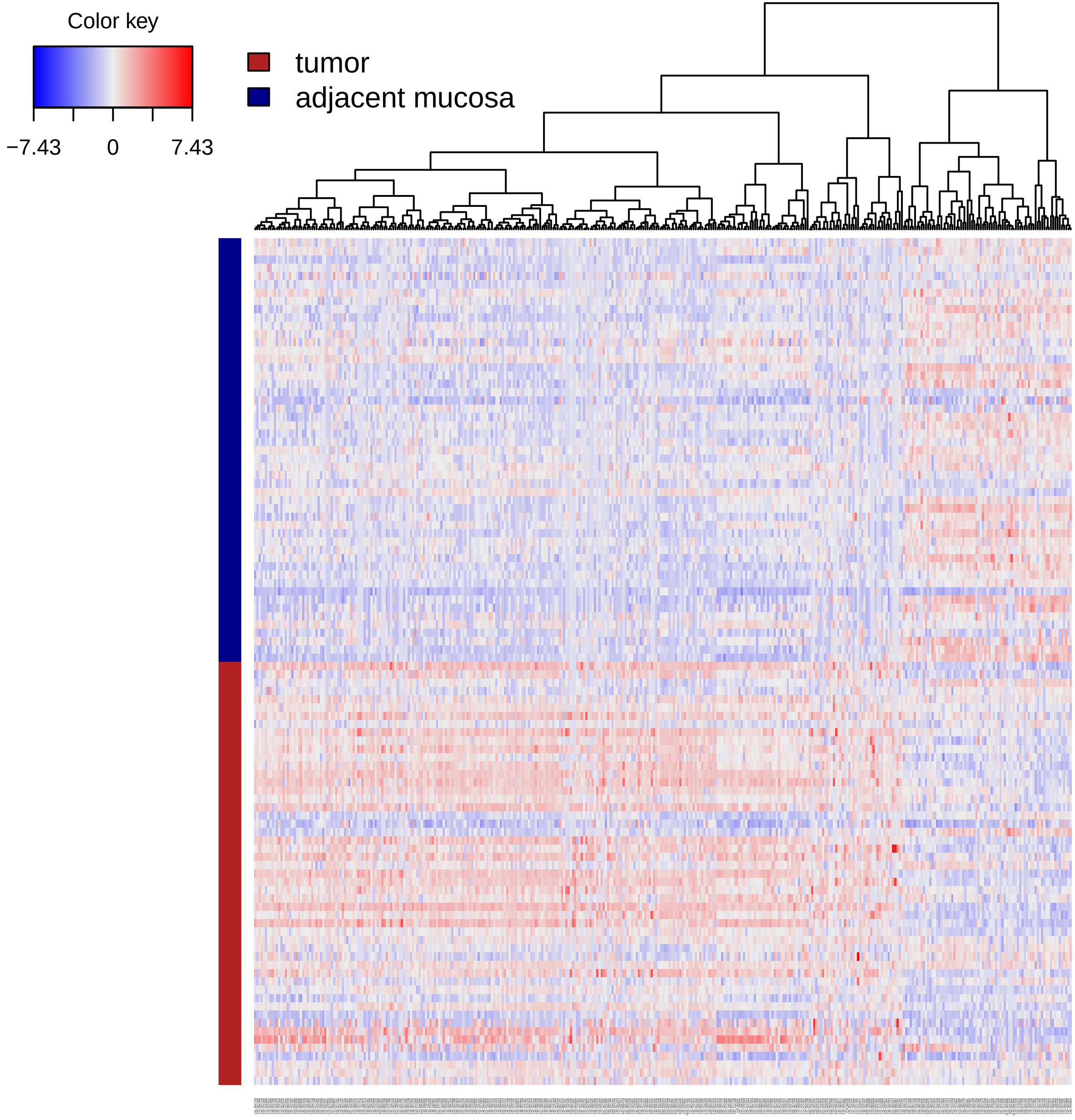


Supplementary Figure 1

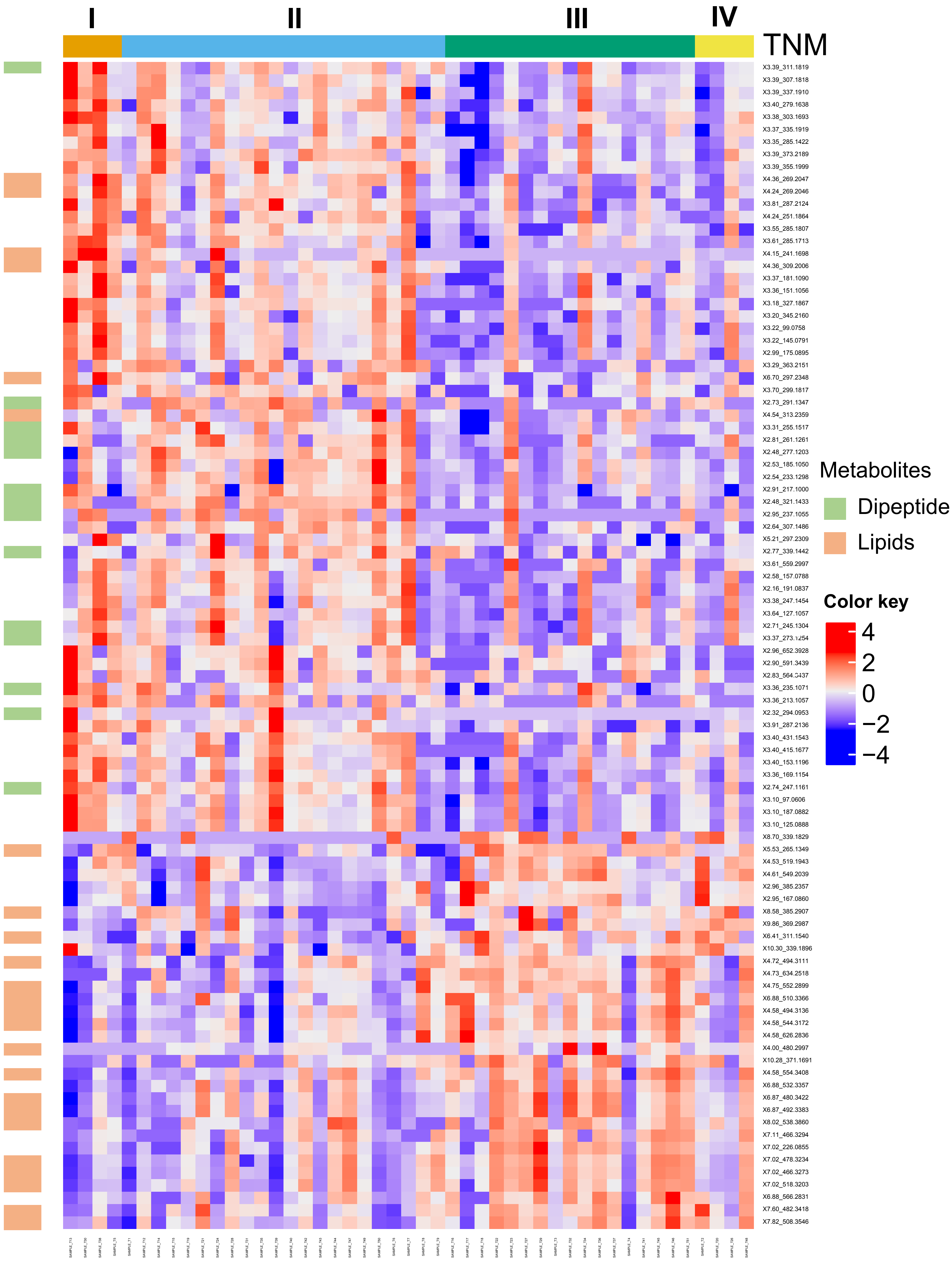


Supplementary Figure 2

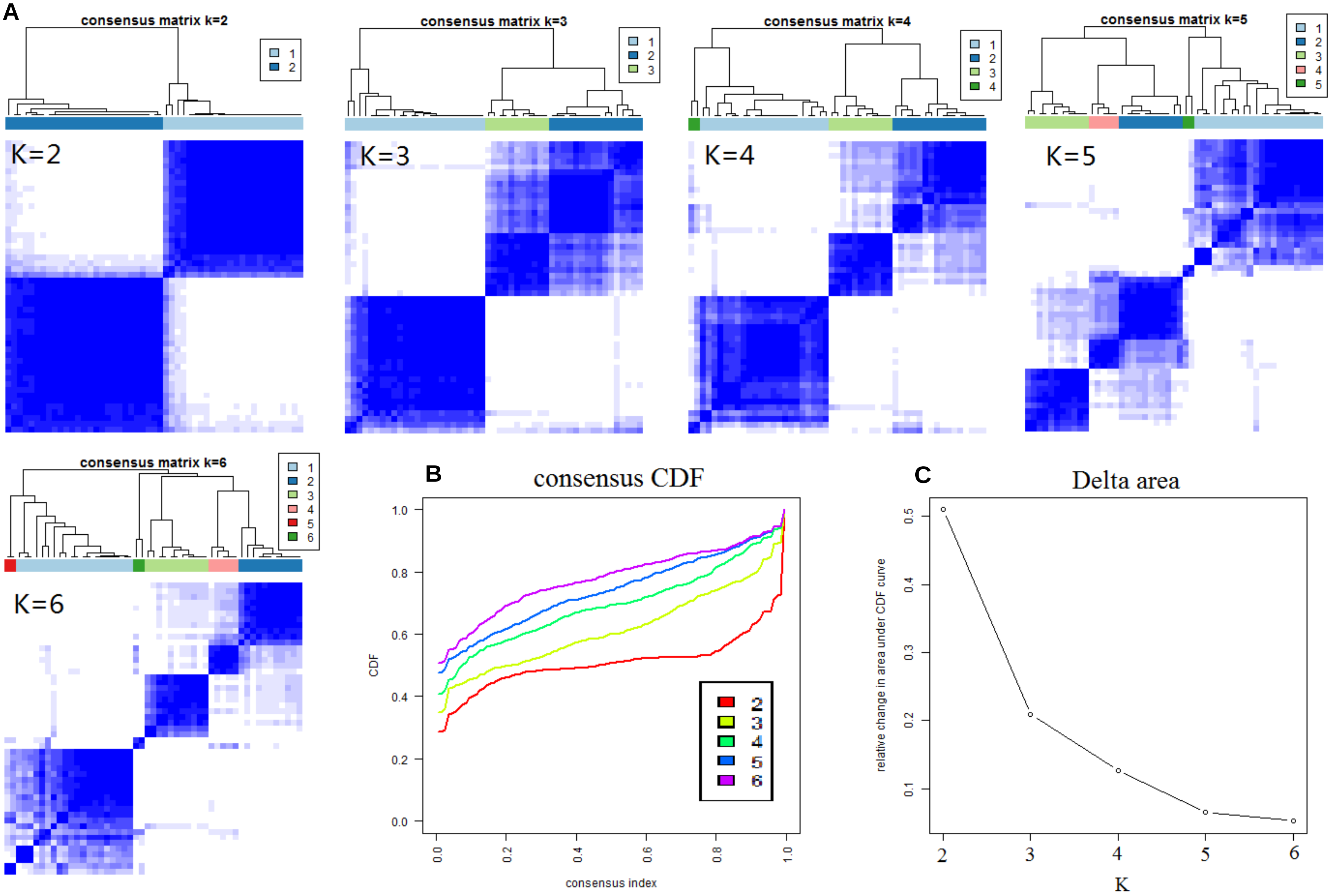


Supplementary Figure 3

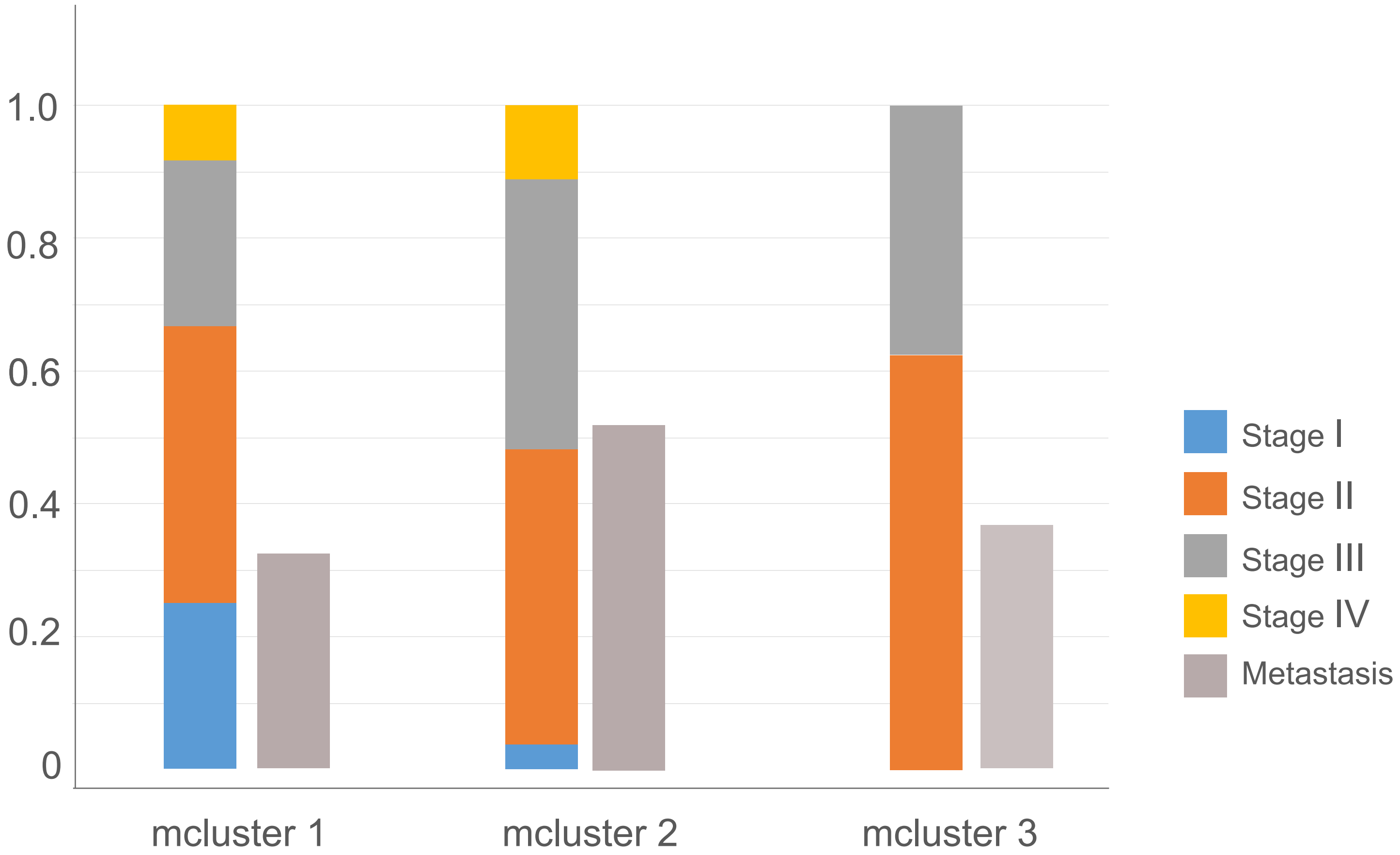


Supplementary Figure 4

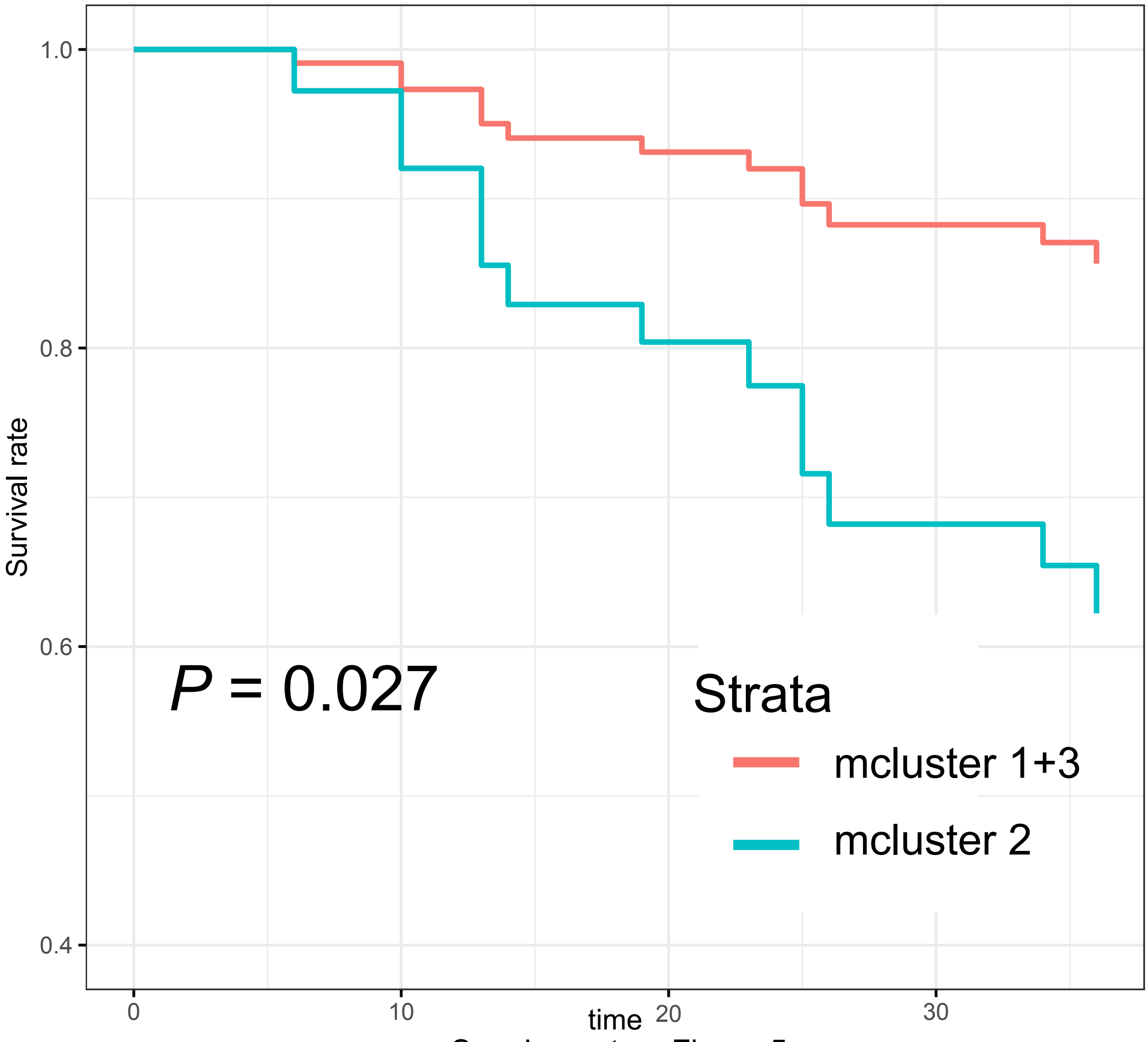


Supplementary Figure 5
